# Zhi-Shi-Wu-Huang attenuates Aβ toxicity in *Caenorhabditis elegans* Alzheimer’s disease models via modulating insulin DAF-16 signaling pathway

**DOI:** 10.64898/2026.03.06.709794

**Authors:** Fahim Muhammad, Yan Liu, Ren Hui, Hongyu Li

**Author notes:** **Correspondence**: Hong-Yu Li, College of Life Sciences, Lanzhou University, Lanzhou, 730000, China.

## Abstract

Alzheimer’s disease (AD) is a common neurodegenerative disorder primarily caused by Amyloid-beta (Aβ) toxicity. Therefore, there is an urgent need to develop novel, effective, and safe drugs to treat AD. Traditional Chinese Medicine (TCM) has a long history of use in protecting against memory impairments. Recently, TCM has attracted growing attention from researchers as a source of potent neuroprotective compounds. In this study, we focus on four TCM herbs with multiple therapeutic properties: *Valeriana jatamansi* (V; 20 mg/mL), *Acori tatarinowii* (A; 10 mg/mL), *Fructus Schisandrae* (F; 5 mg/mL), and *Scutellaria baicalensis* (S; 2.5 mg/mL). The aim is to develop a neuroprotective anti-AD formulation, named “Zhi-Shi-Wu-Huang” derived from V, A, F, and S, and evaluate its efficacy in transgenic *Caenorhabditis elegans* models of AD. These four TCM herbs are among the most potent activators of the HSP-70 promoter, promoting the expression of heat shock protein 70 (HSP-70), which helps prevent protein misfolding and aggregation. Additionally, V, A, F, S, and the Zhi-Shi-Wu-Huang formula were found to reduce reactive oxygen species (ROS) production and enhance the expression of superoxide dismutase-3 (sod-3) and chymotrypsin-like proteasomes. Our findings demonstrate that both the individual extracts (V, A, F, S) and the Zhi-Shi-Wu-Huang formulation significantly reduce Aβ-induced toxicity in transgenic worms by activating the insulin/DAF-16 signaling pathway.

## 1 Introduction

Alzheimer’s disease (AD) is the first most age-related neurological disorder, characterized by neurofibrillary tangles and beta-amyloid (Aβ) accumulated toxicity. AD destroys synapses, induces brain inflammations, and eventually leads to neuronal death with severe brain stroke (1). Many factors are involved in AD pathogenesis, such as Aβ aggregative toxicity (2), tau protein hyperphosphorylations (3), acetylcholine deficiencies (4), inflammation-induced neurodegenerations (5), oxidative stress (6). Among them, the Aβ toxicity postulates broadly accepted, which shows that Aβ aggregations induce inflammations in the brain, synapses impairment, and ultimately lead to the death of neuronal cells. It is so complicated that the pathogenesis of AD has not yet been fully understood. Thus, to decrease Aβ generations, aggregations or accumulations in patients’ brains become the promising anti-AD strategy. Therefore, abnormal Aβ aggregative toxicity clearance is beneficial to neuron protection via stopping the AD-like symptoms. Recently, only a few FDA-approved anti-AD drugs (Memantine, Galantamine, Rivastigmine, and Donepezil) are commercially used but do not significantly affect the patients’ health recovery. These FDA drugs can only delay the onset of dementia, but their prolonged use showed adverse effects comprised gastrointestinal tract, diarrhoea, nausea, vomiting, rapid eye movements, and sleep behaviour disorder (7).

Therefore, new therapeutic approaches are required to discover a plant-based neuroprotective drug to treat or alleviate AD. Traditional Chinese medicine (TCM) has been used for thousands of years with high efficacy and low toxicity record. The TCM-derived therapeutics, including herbal decoctions and individual active constituents, have been used in clinics for a long time (8). Heat shock proteins can prevent false folding and protein aggregation re-fold the misfolded protein, especially the heat shock protein HSP70. To screen the HSP70 activators, we preferentially selected 35 single drugs from the collections and statistics of the traditional Chinese prescriptions and proved the benefit of curing via heat shock protein (HSP) promoter activations. Therefore, the top four HSP70 promoter activator TCMs from the above mentioned 35 single drugs were filtered out, which are Valeriana jatamansi Jones (Zhi Zhu Xiang, “V”), Acorus Tatarinowii (Shi Chang Pu, “A”), Fructus Schisandra Chinensis (Wu Wei Zi, “F”), and Scutellaria baicalensis (HuangQin, “S”). Among them, V is the best HSP70 promoter activator which can activate the HSP70 express in pGL3-HSP70 and pRL-TK co-transfected HEK-293T cells as low as 0.4 mg/mL decoction, while others are 3.2 mg/mL (A), 1.2 mg/mL (S), 5.5 mg/mL (F), respectively. (9). Studies have shown that HSP70 plays a vital role in reducing protein accumulation of α-Syn, Aβ, restoring tau protein in vivo balance, reducing oxidative stress, and inhibiting nerve inflammations(10). At the same time, HSP70 chaperone families control apoptosis at the mitochondrial stage and suppress the stress-induced signaling by inhibiting BAX translocation and enhancing the caspase-3, caspase-8, and Bcl-2 expressions (11). The protection of neurons from aggregated misfolded protein during stress relay on HSP70 family. HSP70 proteins have been expressed with Aβ, the main AD cause, inside the intra-neuronal inclusions in AD brains and the substantia nigra pars compacta (SN) (12). So HSP70 can be used as an anti-Aβ target to screen anti-AD drugs in future studies.

Valeriana jatamansi Jones (V), its family (Caprifoliaceae), more than 200 species, and is a high source of terpenoids, flavonoids, and antioxidants (13). Many of them are well known in Chinese and Tibetan medicine as a part of various formulations to treat migraines, insomnia, nervous tensions, and neuralgia (14). Acori Tatarinowii Rhizoma (A) is a well-known TCM that treats mental disorders (15). A’s active role comprises waking the mind for improving wisdom, dementia, short-term memory issues and treating several other disorders like drive-out appetite, improving pronunciations, forgetfulness, and longevity. The active ingredients comprise flavonoids and antioxidants (16). Fructus Schisandra Chinensis (F) is an active ingredient extracted from the fruit of Schisandra Chinensis Baill. They are rich in flavonoids like Flavin, Glucoside and are widely used as a traditional medicine in China and Eastern Europe (17). F exhibited various pharmacological properties to stop oxidative activities and reduce cellular apoptosis (18, 19). Scutellaria baicalensis Georgi (S) is a perennial herb distributed at a large scale in Eastern Asia (20, 21). S is the most prevalent and multi-purpose herb for neuroprotection, traditionally used in China and other oriental countries (22). S roots are rich in flavonoid contents, including wogonin and baicalein (23). Baicalein, the main flavone in S, strongly inhibited the Aβ toxicity (24). Every drug in V, A, F, S had a long history of traditional use in different regions of China against emotional stresses, nervous disorders, insomnia, epilepsy, memory impairments, and other human diseases. Therefore, in our current study, the four HSP70 promoter activator TCMs combined at a specific ratio (8:4:2:1) (**Table: 1),** named “Zhi-Shi-Wu-Huang” was treated as anti-Aβ toxicity drugs in transgenic *Caenorhabditis elegans* models of AD.

*C. elegans* have an easy culture method, a short life cycle with a simple neurons network, and a conserved nervous system pathway (25). Transgenic *C. elegans* directly express human Aβ_1-42_ peptides in the organism to construct AD-like pathological models. These transgenic AD models can be correctly used to evaluate the Zhi-Shi-Wu-Huang Tang pharmacological properties to eliminate abnormal Aβ. We observed that the treatment of Zhi-Shi-Wu-Huang Tang significantly reduced the Aβ toxicity by paralysis and 5-HT hypersensitivity assays. Besides, the Zhi-Shi-Wu-Huang Tang reduced aggregated Aβ deposits in CL2006 transgenic worms. This anti-Aβ toxicity properties of Zhi-Shi-Wu-Huang formaulation are due to the Insulin/IGF-1 signaling pathway confirmed by DAF-16 RNAi and HSP-16.2 study. Further, Zhi-Shi-Wu-Huang formulation reduced ROS internal cells and enhanced *SOD*-3 expressions. These antioxidant-like properties of Zhi-Shi-Wu-Huang might be due to following the oxidative pathway, but further investigations will proceed in our future projects.

## 2 Material and Methods

### 2.1 Preparation of herbal extract

Weigh 4 selected TCMs (Valeriana jatamansi Jones “V,” Acorus Tatarinowii “A,” Scutellaria baicalensis “S,” and Fructus Schisandra Chinensis “F”) in powder form and boiled for 30 minutes twice. Separated the solutions from the residue, combined, and then filtered. After freeze-drying, further quality control of powders was performed by HPLC. The stock solutions were stored at −20 °C and diluted by two-fold dilution with distilled water to make multiple dilutions for experimental work. To begin with, these drugs were tested by orthogonal experiment on transgenic *C.elegans* AD model CL4176 to find out the best combination of Zhi-Shi-Wu-Huang formulation (named **F-1** in this study). How to calculate this formula orthogonally from four TCM (V, A, S, F) are given in supplementary data file.

### 2.2 *C. elegans* culture and maintenance

*C. elegans* strains used in this study are mentioned in **Table: 2** below. All strains were purchased from the Caenorhabditis Genetics Center, USA, Minnesota. They were maintained by culturing solid nematode growth media (26) and fed with OP50 (*Escherichia coli strains)* as a food source as previously described. To prepare age-synchronized worms, the gravid adults were treated with hypochlorite and M-9 for at least 5-6 min, and later eggs were washed with M9 buffer three times before transfer on fresh NGM plates.

### 2.3 Food clearance test

A food clearance test of *C. elegans* (wild-type N_2_ and transgenic CL4176 (myo-3p:: Aβ_1-42_::let-851 3’UTR) was performed according to a previously described method (27). With brief modifications to determine the suitable, non-toxic concentration of V, A, F, S drug extracts for neuroprotective treatments. OP50 grew overnight and then suspended at a final optical density (OD) of 0.6 in nematode S-basal solution. Every single drug was diluted at 2.5 mg/mL, 5.0 mg/mL, 10 mg/mL, 20 mg/mL, and 30 mg/mL into the *E. coli* suspension to attain the desired nontoxic concentrations. Each well of a 96-well plate received 60 µL of the *E. coli* suspension. Randomly selected 40 synchronized L-1 worms in 10 µL of S-medium were added to an *E. coli* suspension containing a series of V, A, F, S concentrations and then incubated at 20 □C. The plate was covered with aluminium foil to prevent evaporations. The OD of the culture at 595 nm was measured once/d for six days, using a SpectraMax M5 Microplate Reader (Molecular Devices, Silicon Valley, CA, USA). Before measuring OD, each plate was shaken for 10 sec. The fraction of worms alive per well-observed microscopically based on size. The same procedure has been done for the F-1 formula group.

### 2.4 Paralysis assay

CL4176 (myo-3p::Aβ_1-42_::let-851 3’UTR) is a temperature-sensitive transgenic *C. elegans* integrated with human *Aβ*_1-42_ genes and express AD-like phenotype of paralysis at 25 ° C. With short modifications in the previously described protocol (28), *C. elegans* were treated with each V, A, F, S drug at 2.5 mg/mL, 5.0 mg/mL, 10 mg/mL and the Zhi-Shi-Wu-Huang Tang. Control groups were treated without drugs. Each plate was seeded with 49–59 synchronized eggs under 16 ° C. When larvae developed to the L3 stage, the culture temperature increased to 25 ° C for 34 h. Then the non-paralyzed rates were monitored every 2 h. Each *C. elegans* was gently touched with a platinum loop to identify paralysis. *C. elegans* were considered paralyzed if they did not move or only moved their heads. In our experiment, at least three independent biological replicates were performed.

### 2.5 Exogenous serotonin sensitivity assay

After minor changes in the previously described method (29), egg-synchronized transgenic worms CL2355 (snb-1/Aβ1-42) and CL2122 (dvIs15) were placed at 20 °C on NGM plates seeded with OP50 for 48 hrs. CL2122 worm strains were used as the positive control (without Aβ expressions). Later both the worms were treated with V, A, F, S drugs at 2.5 mg/mL, 5.0 mg/mL, 10 mg/mL, and the F-1 for another 48 hrs. 30 worms in each group were washed with M-9 buffer three times and were transferred into 96 wells microtiter plate containing 1 mg *Serotonin* (creatinine sulfate salt; *Sigma*) in 200 μL M-9 buffer. After 5 min, we have counted the worm’s paralysis for 5 sec.

### 2.6 Fluorescence staining of oligomer Aβ deposits

Eggs synchronized transgenic worms CL2006 (*unc-54*/human Aβ_1-42_) were maintained and grown on NGM at 20 ° C up to the L-4 stage, as defined previously (30). Then worms were transferred to the NGM plates containing V, A, F, S drug at 2.5, 5.0, and 10 mg/mL and the F-1, incubated for 48 hrs at 20 ° C. With minor modifications in the previously described protocol; we performed Thioflavine-S (Th-S) staining. In short, worms were gathered by washing with M-9 and were fixed in 4% paraformaldehyde in PBS (pH7.4), incubated at 20 ° C for 24 hrs. The fixative solution was replaced by permeabilization solutions (5% fresh β-mercaptoethanol, 1% Triton X-100, 125 mM Tris-HCl, pH7.4) and incubated at 37 °C for another 24 hrs. Later the worms were washed with PBS-T (PBS plus 0.1% Triton X-100) thrice and stained with 0.125% Th-S (*Sigma*) in 50% ethanol for 4-6 min, and the samples were washed out with 50% ethanol thrice for 2-4 mins. Stained samples were re-suspended in 100 μL PBS (pH 7.4), and finally mounted on agar glass pads and observed by fluorescence microscopy (BX53; Olympus, Japan) under the same setting parameters. Th-S positive deposits were quantified in the forepart of the pharyngeal bulb in every worm by Image J software. The Fluorescence of randomly selected worms was statistically observed in each group. At least three independent biological replicates were performed in our experiment. Here wild-type N2 is used as amyloid deposit control.

### 2.7 Western blot

After V, A, F, S individual drug treatment at 10 mg/mL for 3 d and 34 hrs temperature upshift to 25 □C, transgenic CL4176 *C. elegans* was washed with fresh PBS and stored in a □80 refrigerator. Wild-type N2 was used as a negative control group. Similarly, the same procedure was applied to the F-1 to treat *C. elegans*. The next day, whole animal frozen pellets were taken on ice and lysed with lysis buffer (2X NP-40 buffer + 1 mM PMSF) by tissue grinding pestles in 1.5 ml Eppendorf tubes for 40-60 sec, later boiled for 10 min and centrifuge at 14000 g in 4□C. After centrifugation, the supernatant was transferred into a new tube, and the pellet was discarded. The supernatant containing the required protein for performing immunoblot and protein concentration measured by BCA kit. Before loading the protein samples on the 16% Tricin gel for separations, Protein extracts were boiled for 10 min with a loading buffer containing SDS and DTT. After the gel run, proteins and marker bands were transferred to 0.22 µm PVDF membranes. Then blocked the protein bands with 10% skimmed milk in 1X TBST for two hrs. After careful washing, the primary antibody 6E10 (Biolegend, 803,001) at 1:1000 dilution for the whole night at 4 □C refrigerators was used to detect the human Aβ_1-42_. The next day washed the membrane with 1X TBST for 5 minutes thrice and then used a secondary antibody (goat-anti mouse, 1:5,000; BioRad, Hercules, CA, USA) for 1:30 hrs at room temperature. Bands intensity visualized by the horse-radish peroxidase chemiluminescence (ECL Lumi-Light, Roche, Germany) in Vilber Fusion FX gel documentation system, 2019 https://www.vilber.com/fusion-fx/. Later, images were quantified by ImageJ software (http://imagej.net/).

### 2.8 RNA interference (RNAi) Assay

Eggs synchronized transgenic CL4176 (myo-3p::Aβ_1-42_::let-851 3’UTR) worms were maintained on NGM/RNAi plates for two generations as described previously (30). L-1 larvae were fed to RNAi bacteria for DAF-16/SKN-1 RNAi in the first generations. These RNAi NGM plates contained 100 μg/mL Ampicillin, 10 μg/mL Tetracycline, and 1 mM Isopropyl β-D-1-Thiogalactopyranoside (IPTG) and were seeded with *E. coli* HT115 cloned targeted with DAF-16, SKN-1 genes, and empty vector L4440. After two generations of propagations on RNAi NGM plates, synchronized L1 worms were treated with or without V, A, F, S at 10 mg/mL and F-1 and incubated at 16 °C for 72 hrs, then upshifted the temperature to 25 ° C for 28 hrs. The number of paralyzed worms was counted as the method defined in the Paralysis assay.

### 2.9 Subcellular nuclear localization of DAF□16::GFP and SKN-1::GFP assay

Transgenic strain TJ356 (*daf*-16a/b::GFP), integrated with *daf*-16a/b promoter fused to GFP, has been used and confer the expression of daf-16a/b on V, A, F, S at 10 mg/mL and F-1 treatment. With short modifications in the previously described protocol (31), L-1 age-synchronized worms were treated with or without a drug for 72 hrs. The positive control groups were treated at 37 □C for 30 min before observing fluorescence. While, for SKN-1::GFP localization, we used LG333 (*skn*-1b::GFP) transgenic worms and treated with V, A, F, S, and F-1 for 72 hrs at 20 □ C. At least 25 randomly selected worms from each group were observed on a glass slide by fluorescence microscopy. Later, we examined the subcellular distributions of daf-16a/b::GFP and *skn*-1b::GFP by fluorescence microscope and the ratio of the worms with SKN-1 or DAF-16 nuclear localization counted.

### 2.10 The HSP-16.2::GFP and GST-4::GFP expression assay

Transgenic strain TJ375 (*hsp*-16.2::GFP), integrated with hsp-16.2 promoter fused to GFP, have been used and confer the expression of *HSP*-16.2 on V, A, F, S at 10 mg/mL and F-1 treatment. With short modifications in the previously described protocol (31), L-1 age-synchronized worms were treated with or without a drug for 72 hrs. The positive control groups were treated with Juglone at 20 mM for the same period. While for SKN-1, synchronous L-1 larvae of transgenic CL2166 (*gst*-4p::GFP) worms moved onto fresh NGM plates with or without V, A, F, S at 10.0 mg/mL and F-1 for 72 hrs at 20 °C. Positive control worms were treated with 1.5 mM NaAsO2 at 20 °C for 1 h before observing fluorescence. Later, worms were washed with M-9 buffer, mounted onto an agarose pad slide with 20 mM sodium azide, and enclosed with coverslips. Then images were observed under the fluorescence microscope (BX53; Olympus Corp., Tokyo, Japan). Later fluorescence intensity of images was quantified by ImageJ software.

### 2.11 *SOD*-3::GFP reporter assay

The *SOD*-3 (superoxide dismutase) assay for antioxidants expressed previously described was performed (32). The reporter gene assay was conducted using transgenic strains CF1553, strongly expressed *SOD*-3::GFP. L-1 synchronized *C. elegans* were transferred to the NGM solid media plates with or without V, A, F, S at 10 mg/mL and F-1 formula drug incubated at 20 ° C for 72 hrs. Later, *C. elegans* were washed with M-9 buffer, mounted on an agarose pad with 10 μL of 20 mM sodium azide, and closed with a coverslip. Positive control was given heat shock at 37 ° C for 30 min, and fluorescence intensity of the treated group’s images was observed *via* fluorescence microscope (Olympus, BX53, Japan). The quantification of fluorescence intensity of *C. elegans* images was calculated using Image-J software.

### 2.12 Chymotrypsin-like proteasome activity in *C. elegans*

Chymotrypsin-like activities of proteasomes in transgenic CL4176 (myo-3p::Aβ_1-42_::let-851 3’UTR) worms were examined using a previously defined protocol (33). Worms were treated with V, A, F, S at 10 mg/mL and F-1. Control groups were treated without drugs. With brief modifications, *C. elegans* were lysed in proteasomes lysis solution (NP-40+1mM PMSF) via tissue grinding pestles for homogenization. Later, centrifuged the lysate at 14,000 g for 10 min at 4□C, then transferred the supernatant into a separate 1.5 mL Eppendorf tube containing the protein plus proteasome subunits, and discarded the pellet. 20 μL from the supernatant was loaded on a 96-well plate containing a 30 μL fluorogenic substrate for each test. LLVY-R110-AMC (Sigma-Aldrich, MAK172-IKT, USA) was used as the fluorogenic substrate to examine the ubiquitin-like proteasome activities in the worm’s lysate. Proteasome cleavages the substrate LLVY-R110 and generates a green fluorescence signal on R110 separations. The lysate and fluorogenic substrate were incubated in a black 96 wells microtiter plate for 1.5 h at 37 □C. Then, reading was taken at 490 nm excitation wavelengths and 525 nm emission wavelengths at 10 min intervals for 1 hour by M2 Microplate Reader SpectraMax (Molecular Devices, Silicon Valley, CA, USA).

### 2.13 Intracellular ROS

We used a previously described protocol (34) with brief modifications to measure the antioxidative properties of the drugs V, A, F, S individually and F-1. Therefore, we performed an H2DCF-DA assay at 50 μM in wild-type N2 and transgenic CL4176 (myo-3p::Aβ1-42::let-851 3’UTR) worms. Both wild-type N2 and transgenic CL4176 C. elegans were treated with or without V, A, F, S at 10 mg/mL and F-1 for 72 h at 20 □ C. After that, washed the worms with M9 and transferred (at least 30) worms to 96 wells microtiter plate with 50 μM of H_2_ DCFDA in 200 μL PBS per well. Later wild-type N2 worms were washed and transferred to the fresh NGM plates for 30-40 min crawling and then observed the fluorescence intensities of each treated worms under the BX53 fluorescence microscope. While for transgenic CL4176, worm’s fluorescence was measured immediately at 20 min intervals for 200 min, using excitation 485 nm and emission at 520 nm by a SpectraMax M2 Microplate Reader.

### 2.14 Lifespan

AD is an age-related disorder. Therefore, we have performed a lifespan test with a simple modification in the previously described method (35) to examine the effects of the V, A, F, S drugs on worm age. Both wild-type N2 and transgenic CL4176 were treated with V, A, F, S at 10.0 mg/mL and F-1 for 72 hrs at 20 □ C. Control groups were treated without drugs. Later, washed with M-9 and transferred into fresh NGM plates seeded with *E. coli* OP50 strains and incubated at 20 □C until all worms became dead. Each worm was marked as live, dead, or censor at day intervals, considered by their pharyngeal pumping. The worms that perform normal behaviour or have their pharynges pump were considered live. The worms scored dead based on the pharyngeal pump stopped working by gentle touch. Besides, the worm exhibiting abnormal behaviours, including unusual death, or crawling out of the plate, was considered a censor and excluded from the calculation. Nevertheless, the observation was conducted until all the worms became dead and the lifespan compared to the control.

## 3 Results

### 3.1 Zhi-Shi-Wu-Huang’s food clearance assay

For details, please refer to the supplementaory figure **S03.**

### 3.2 Zhi-Shi-Wu-Huang recovered AD-like symptoms prompted by Aβ1-42 accumulated toxicity

The CL4176 (myo-3p::Aβ_1-42_::let-851 3’UTR) is the transgenic worm, express tissue-specifically human Aβ(_1-42_) in muscle, which is induced by temperature raised to 25 □C. With the expressions and the accumulations of Aβ deposits into the worm’s muscles and shows severe paralysis phenotype. Therefore, we treated the CL4176 worms with V, A, F, S at 2.5 mg/mL, 5.0 mg/mL, 10 mg/mL, and F-1 to validate the anti-Aβ toxicity effects. All V, A, F, and S concentrations significantly delayed worms’ paralysis process **Fig. 2**. Similarly, we have treated the F-1 in the same way. We observed F-1 had the best excellent therapeutically effect on Aβ toxicity reductions than single drugs on treatment.

**Figure (2):**
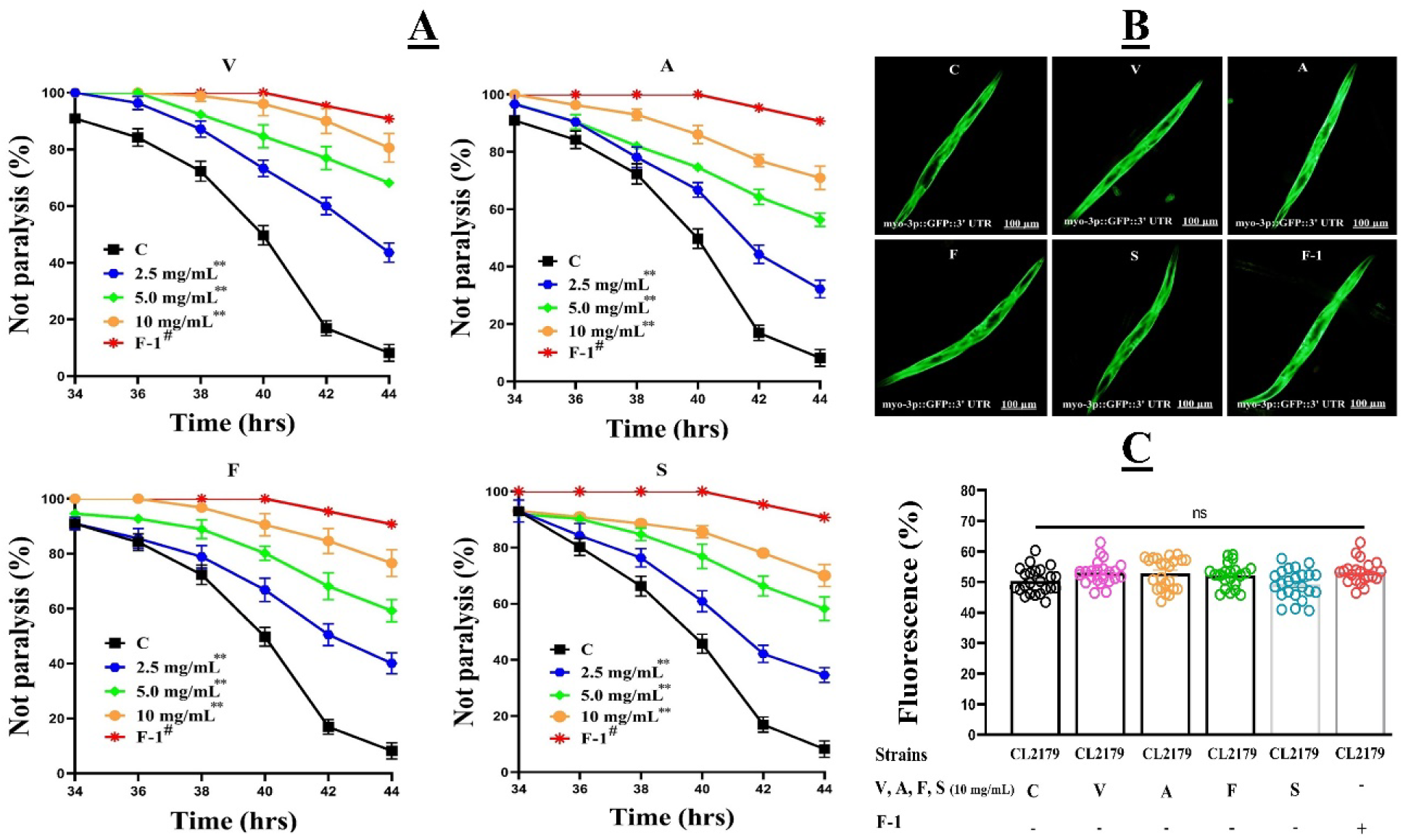
(A) V, A, F, S, and F-1 together significantly reduced the human Aβ (myo-3p::Aβ(_1-42_)::let-851 3’UTR) accumulated toxicity and delayed AD-like symptoms in the CL4176 transgenic worms. Paralysis was observed in each V, A, F, S treated at 2.5 mg/mL, 5.0 mg/mL, 10 mg/mL, and untreated groups for 34 hrs of temperature upshift to 25 C. The same procedure was done with the F-1 formula drug. Data calculated by mean±SD, n=4. ANOVA one-way followed by Tukey’s test was used to calculate the significant difference. ** indicated the significant difference between drugs V, A, F, S treated and control groups *p*≤0.005 (**). # showed the significant difference between F-1 and control groups *p*≤0.005. (B) Fluorescence images of myo-3::GFP expression in CL2179 worms. CL2179 is used here as an Aβ transcription control. The scale bar is 100 μm; (C), indicating each treated group’s quantitative analysis of myo-3::GFP expressions intensity. Data were calculated by mean ± SD, n = 30. ns representing no significant

However, to evaluate whether V, A, F, S, and F-1 could non-specifically work any exogenous protein (Aβ) expressions or exclusively act on Aβ toxicity, CL2179 [*myo-3p*::GFP: 3′UTR (36) + *rol-6* (su1006)] was used as transgenic background control. We treated V, A, F, S at best effective concentrations 10 mg/mL and F-1 with CL2179 worms for the same period with paralysis assay worms. Expectedly, the GFP fluorescence intensity of treated groups revealed no significant difference found in **Fig. 2 (B&C)**. Therefore, we can conclude that V, A, F, S, and F-1 significantly eliminated the Aβ toxicity in muscle cells and alleviated AD-like symptoms in the treated worms *p*≤0.005 (**).

### 3.3 Zhi-Shi-Wu-Huang suppresses neuronal Aβ1-42 expressions

The paralysis phenotype in the muscle of the Aβ (myo-3p::Aβ_1-42_::let-851 3’UTR) expression strain CL4176, used above, has been established and used to illustrate several important molecular processes related to Aβ toxicity. To relate Aβ toxicity with neuronal functions, we characterized a behavioural phenotype of the transgenic strain CL2355, in which Aβ expressed in the neuronal region. For that, we selected a neuronal controlled behaviour assay 5-HT sensitivity in CL2355 and CL2122 worms. CL2122 is without Aβ expression and is used as control strains of CL2355. CL2355 are integrated with human Aβ_1-42_ peptides in pan neurons, which showed hypersensitivity response on exposure to a chemical Serotonin (5-HT) (37). 5-HT is an important neurotransmitter that controls the worm’s egg-laying, smell sense learning, mating, and movement behaviours. Exogenous 5-HT can stop the worm’s movement, leading active worms into paralysis. Similarly, *C. elegans* strain CL2355 shows 5-HT hypersensitivity when Aβ is over-expressed in the nerve system. V, A, F, S at 2.5 mg/mL, 5.0 mg/mL, 10 mg/mL and F-1 significantly alleviated exogenous 5-HT hypersensitivity tempted by Aβ toxicity. But no significant difference among the same treatment groups was observed in CL2122 strains, the transgenic background control of CL2355 **Fig. 3** (V, A, F, S). Therefore, it concluded that V, A, F, S, and F-1 could exclusively stop AD-like symptoms induced by Aβ toxicity in muscle cells and neurons. Our experimental assessment showed combined drugs in the formula “F-1” responded better than individual drugs to 5-HT hypersensitivity induced by Aβ toxicity.

**Figure (3):**
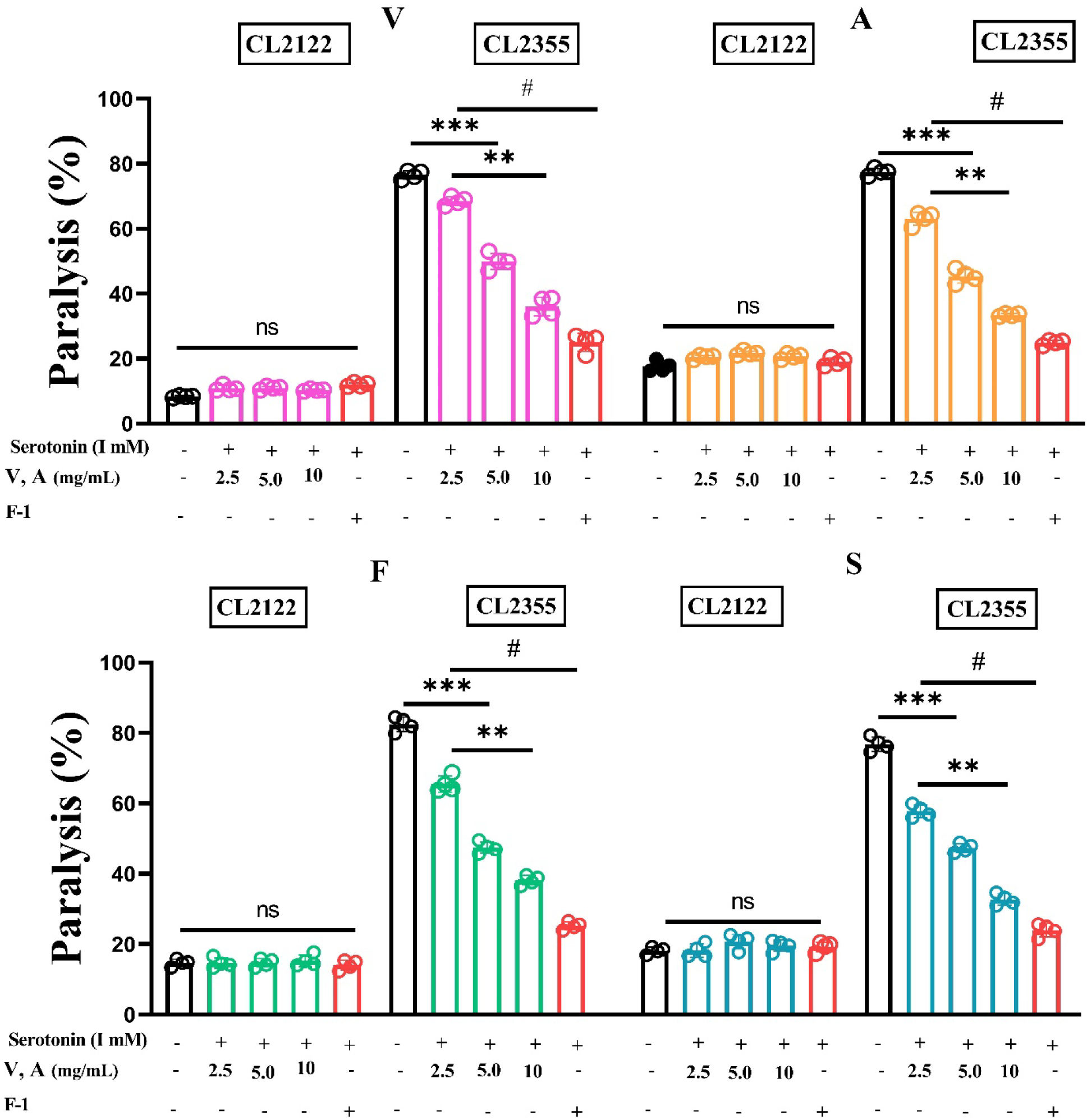
Explaining the effect of V, A, F, S on the exogenous 5-HT hypersensitivity in snb-1:Aβ_1-42_ worms. Data were calculated by mean ± SD, n = 3 (40-60 individuals per group). n representing the no significant difference was observed in CL2122 worms treated with or without V, A, F, S. ** showing the significant difference between V, A, F, S treated at 2.5 mg/mL, 5.0 mg/mL, 10 mg/mL and untreated groups. While # representing the significant difference between F-1 and control group in transgenic strains CL2355 (*p*≤0.005) via one-way ANOVA, followed by Tukey’s test).

### 3.4 Zhi-Shi-Wu-Huang reduced Aβ1-42 positive geposits in transgenic *C. elegans*

To explore the Zhi-Shi-Wu-Huang formula effects on the aggregations of the large Aβ species in the form of Aβ deposits in the worm’s muscles, we selected CL2006, a transgenic *C. elegans* (unc-54/human Aβ_1-42_) that consistently overexpresses human Aβ_1-42_. After V, S, F, S, and F-1 treatment, Aβ deposits in each worms’ forepart of the pharyngeal bulb were calculated after Th-S staining (28). We found the V, A, F, S at 2.5 mg/mL, 5.0 mg/mL, 10 mg/mL, and F-1 on treatment significantly decreases the aggregations of Aβ_1-42_ deposits, in dosed manner **Fig. 4 (A&B)**. Memantine is used here as a positive control group. Memantine is a selected NMDA receptor antagonist, and evidence indicates that it lowers the Aβ level, decreases Aβ plaques, and significantly reduces Aβ deposits (38) **Fig. 4 (A&B)**.

**Figure (4):**
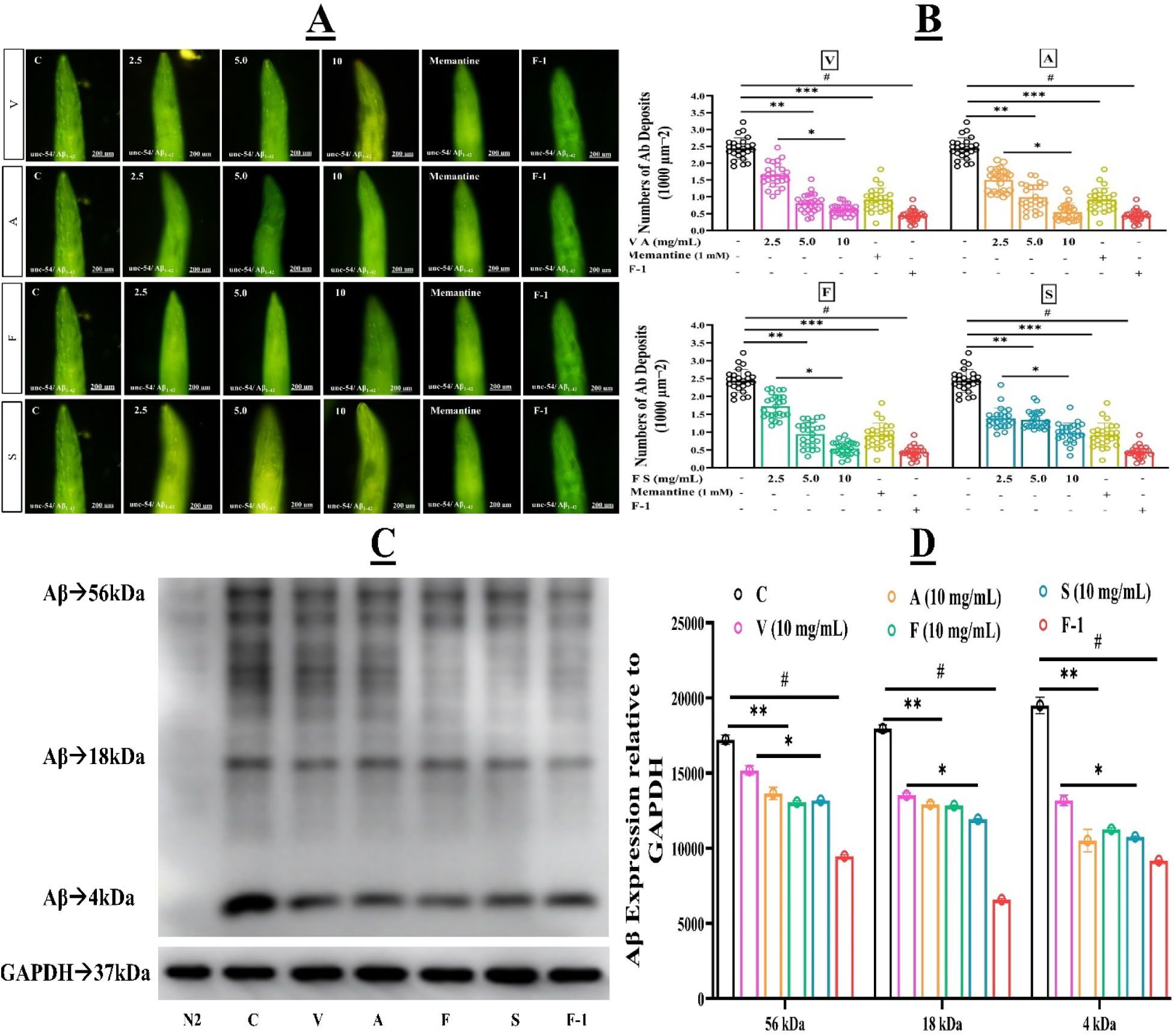
**(A)** Aβ positive deposits stained by Th-S in CL2006 _(_*unc-54*/Aβ_1-42)_ worms after V, A, F, S at 2.5 mg/mL, 5.0 mg/mL, 10 mg/mL and F-1 treatment. **(B)** The quantitative analysis of Aβ positive deposits of each treated and untreated group after ThS staining. Data were calculated by mean ± SD, n = 3 (25 individuals per group). Memantine is used here as a positive control. ** shows a significant difference between treated and respective control (*p*≤0.005), *** denotes the significant difference between Memantine and untreated groups (*p*≤0.005). At the same time, # represents the significant difference between F-1 treated groups vs control groups (*p*≤0.005). The significant difference was calculated by ANOVA (one-way), followed by Tukey’s test. **(C)** V, A, F, S reduced the Aβ protein accumulated expressions by the immunoblot. Results showed that V, A, F, S drug groups significantly reduced the Aβ accumulations in CL4176 worms than untreated groups. In combination, the F-1 showed improved results in reducing Aβ aggregative toxicity. **(D)** Graphical effects of immunoblot band intensities of V, A, F, S in single and F-1 relative to the GAPDH (37 kDa). ** shows the significant difference of (*p*≤0.005) between V, A, F, S treated and untreated groups. # indicated the substantial difference in (*p*≤0.005) between F-1 formula drug and untreated groups.

### 3.5 Zhi-Shi-Wu-Huang reduced Aβ1-42 accumulated toxicity

Western blotting was performed to identify the Aβ_1-42_ protein accumulations. To further verify the Zhi-Shi-Wu-Huang Tang effects in reducing protein Aβ_1-42_ accumulated toxicity in transgenic *C. elegans*. We have treated CL4176 worms with V, A, S, F at 10 mg/mL, and F-1 conducted the western blotting using antibodies (primary and secondary) after temperature upshift. Besides, we also treated CL4176 worms with F-1 implemented the same protocol. GAPDH (37 kDa) was used as a positive control, and data were measured with GAPDH protein expression relative to drug-treated groups. Western blot results proved that drugs groups V, A, F, S on the treatment reduced Aβ_1-42_ accumulated expressions compared to the untreated groups. In contrast, F-1 showed significantly better results in lowering Aβ_1-42_ accumulated toxicity about 34.7 %, *p*≤0.005 in CL4176 transgenic worms on treatment than untreated groups **Fig. 4 (C&D)**.

### 3.6 Zhi-Shi-Wu-Huang reduced AD-like symptoms in transgenic worms interceded by DAF-16 but not SKN-1 pathway

DAF-16 is a crucial component of the Insulin/IGF-1 signaling pathway to regulate many gene expressions for defending against various stresses, including oxidative stress and proteotoxicity. DAF-2 pathway activations have been beneficial for delaying paralysis induced by human Aβ accumulated toxicity in the transgenic strains CL4176. To examine whether V, A, F, S required DAF-16 via following the DAF-2 insulin-like pathway, we treated the worms at a highly effective dilution of 10 mg/mL to protect the worms from severe stress of abnormal Aβ generations and its accumulations. For that, we have knocked down the expression of the transcription factor DAF-16 by RNAi in transgenic worms CL4176 and found that the inhibitory effect of V, A, F, S at 10 mg/mL on worm paralysis significantly decreased **Fig. 5 (A&B)**. These results indicated that V, A, F, S reduced Aβ accumulated toxicity through DAF-16-based mechanisms.

**Figure (5):**
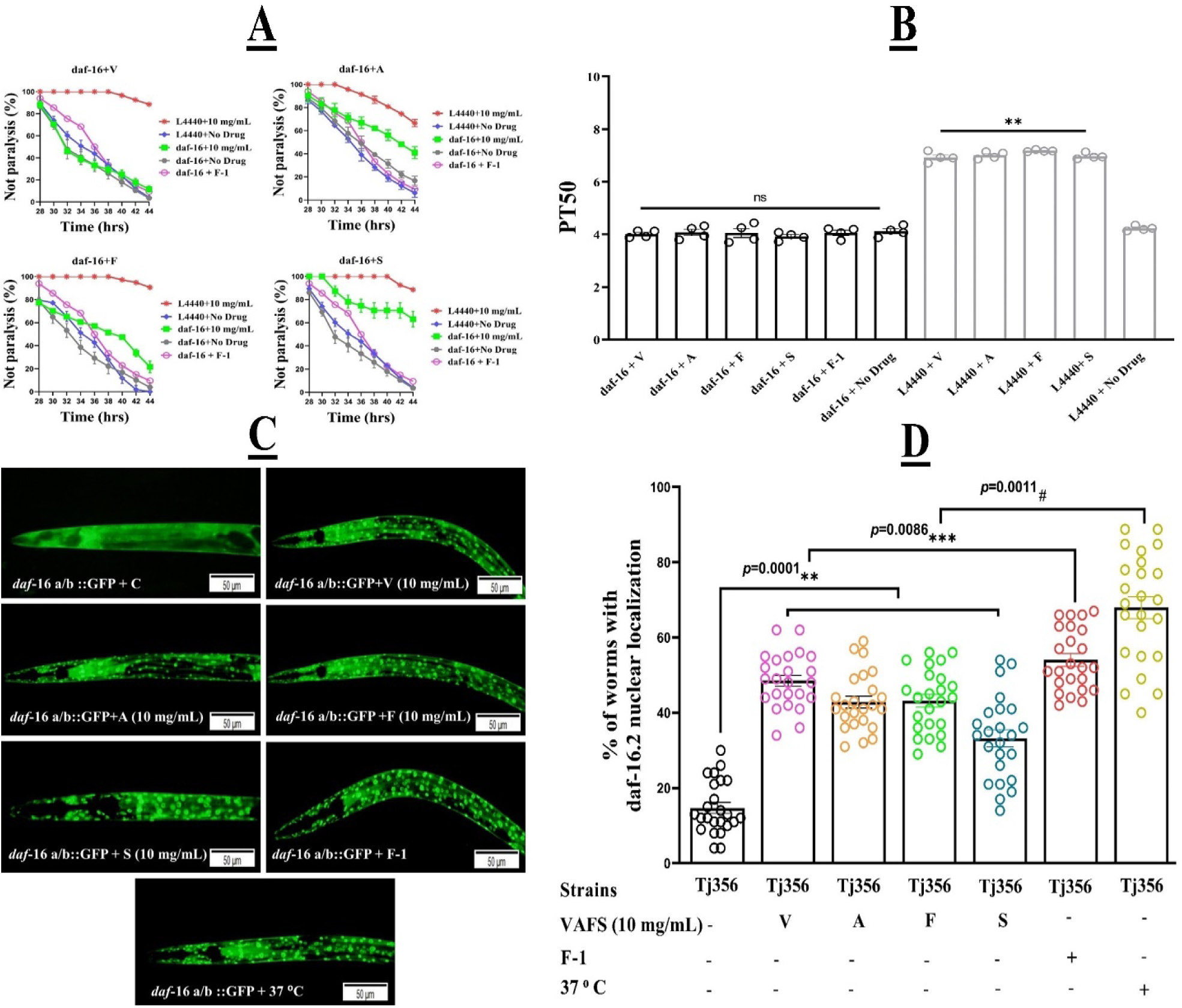
(A&B). V, A, F, S on the treatment inhibiting worm AD-like symptom was independent of DAF-16 Insuline/IGF signalling pathway. **(A)** Fractions of not paralysis curves in CL4176(myo-3p::Aβ_1-42_::3′UTR (long) worms after treatment with or without daf-16 RNAi; (B) PT50s of each treated group in part A, there is a significant difference among these groups when letters are different **(*p*≤0.005). While n represents no significant difference between daf-16 RNAi/ V, A, F, S treated and untreated groups. **(C)** V, A, F, S and F-1 effects on nuclear localization of DAF-16 in TJ356 (daf-16a/b::GFP) worms. Here worms treated at 37 °C for 30 min were considered the positive control group. The scale bar is 50 μm. **(D)** Quantitative analysis of each treated group by ImageJ software. ** denotes the significant difference among V, A, F, S treated groups and untreated group (*p*≤0.005), *** indicating the significant difference between F-1 and untreated group (*p*≤0.005), while # showing the significant difference between the positive control group and untreated group.

Transgenic worms TJ356 (*daf*-16a/b::GFP) carries a GFP fusion protein DAF-16::GFP; thus, DAF-16 activations can be markedly visualized by GFP nuclear distributions. Therefore, in the present work, we treated the TJ356 worms with V, A, F, S at 10 mg/mL and F-1. We observed the nuclear localization of DAF-16 in the tested worms **Fig. 6 (C&D)**. These results indicated that DAF-16 was required for V, A, F, S in resisting the Aβ toxicity in treated worms.

**Fig. 6.**
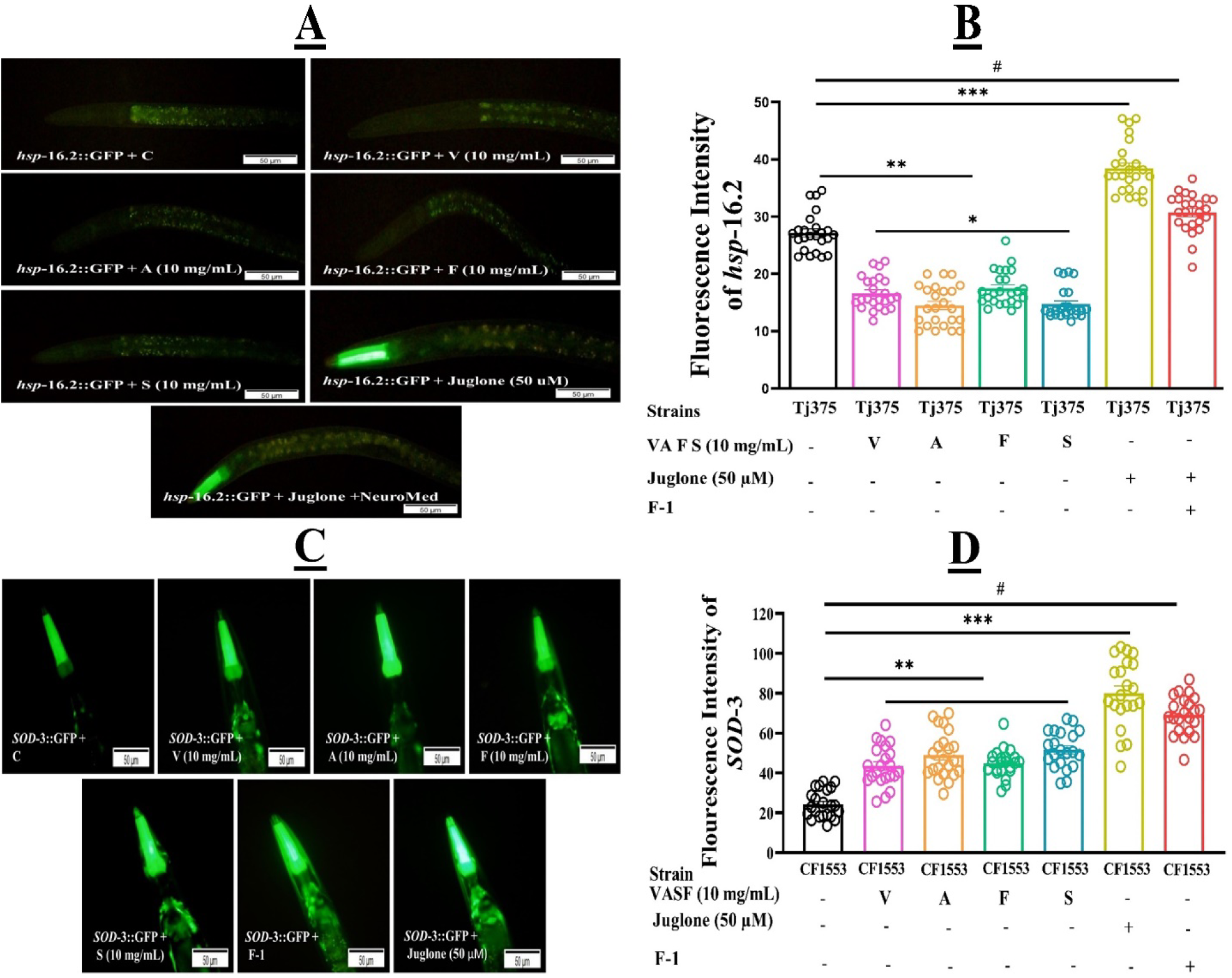
**(A)** V, A, F, S and F-1 effects on nuclear localization of DAF-16 in Tj375 (*hsp*-16.2::GFP) worms. Here worms treated at 50 µM Juglone for 1 hr are considered the positive control group. The scale bar is 50 μm. **(B)** Quantitative analysis of each treated group by image software. ** denotes the significant difference among V, A, F, S treated groups and untreated group (*p*≤0.005), *** indicating the considerable difference between F-1 and untreated group (*p*≤0.005), while # showing the significant difference between the positive control group and untreated group. **(C&D)** Reporter gene *SOD*-3::GFP increased the antioxidant enzymes in the CF1553 *C. elegans* on V, A, F, S, and F-1 treatment. (A) Clearly showing the GFP enhanced expressions of *SOD*-3 in the worms treated with V, A, S, F at 10 mg/mL compared to untreated groups. **(D)** Presenting the graphical form of *SOD*-3 reporter gene expressions results, showing that V, A, F, S improved the antioxidative enzyme actions while alleviating oxidative stress, a significant cause of PD. Data were calculated by mean+SD. Juglone treatment at 50 µM to the worms was taken as the positive control. ** describing the substantial difference in *p*≤0.005 between V, A, F, S treated and untreated groups. *** representing the significant difference of *p*≤0.005 between F-1 and control group. Similarly, # means the significance difference of p≤0.005 between the positive control and untreated groups.

Similarly, HSPs are Heat Shock Proteins with Low-molecular-weight (12-43 kDa). In the case of thermal and oxidative stresses, HSP proteins responded to neutralize the anxiety. Furthermore, HSPs also prevented the accumulated toxicity of numerous proteins such as Aβ. HSP16.2 can suppress Aβ toxicity by helping abnormal protein degradations or refolding and sequestration in AD-treated models. For that, we have selected transgenic models Tj375 (*hsp*-16.2p::GFP) with the *hsp*-16.2::GFP promoter and treated with V, A, F, S drugs at 10 mg/mL and the F-1. Here, V, A, F, S, and F-1 treatment did not enhance the expression of *hsp*-16.2::GFP, representing the reductions in *Aβ* toxicity after V, A, F, S treatment was not mediated by molecular protein HSP-16.2. Then we exposed the transgenic worms Tj375 with Juglone at 50 µM for one hrs. Juglone is an oxidative stress inducer used as a positive control group. Later we treated the Juglone exposed worms with V, A, F, S, and F-1. However, V, A, F, S, and F-1 decreased the up-regulation of hsp-16.2 expressions induced by Juglone **Fig. 6 (A&B)**. It confirmed that V, A, F, S, and F-1 could reduce Aβ toxicity by suppressing oxidative stress via enhancing the antioxidant activities. However, further research on V, A, F, S, and F-1 antioxidative pathways is needed.

As we know, AD is an age-related neurodegenerative disorder. The generation of mitochondrial ROS via oxidative stress is a significant risk factor that leads to abnormal Aβ toxicity. Sodium superoxide dismutase (*SOD*) is an antioxidant enzyme that reduces oxidative stress and decreases the AD-like symptoms via preventing the generations of abnormal Aβ and its accumulation. *SOD*-3 locates downstream of DAF-16 in CF1553 worms. As we know, DAF-16 is a transcription factor and plays a crucial component in the DAF-2/DAF-16 insulin-like signaling pathway. The activation of DAF-16 will lead to elevated expression of *SOD*-3 genes in responses to a wide range of stressors **Fig. 6 (C&D)**. Therefore, our results supported that V, A, F, S, and F-1 reduced oxidative stress by enhancing the *SOD*-3 expressions and attenuated the Aβ toxicity via the Insulin/IGF activations like the DAF-16 pathway.

Like the Nrf2 pathway in mammals, In *C. elegans*, we have the SKN-1 pathway. SKN-1 transcription factor associated with the Cap’n’collar family regulates phase II detoxification response genes to take a protective effect. To identify whether V, A, F, S were required to hinder the toxicity induced by abnormal Aβ generations and its aggregations, the skn-1 expression was knocked down by RNAi. The result showed that skn-1 RNAi did not counteract the V, A, F, S inhibitory effect on worm paralysis **Fig. 6**. It suggested that SKN-1 was not required for V, A, F, S drugs to ameliorate AD-like symptoms of the worm paralysis phenotype induced by Aβ toxicity **Fig. 7 (A&B)**.

**Fig. 7.**
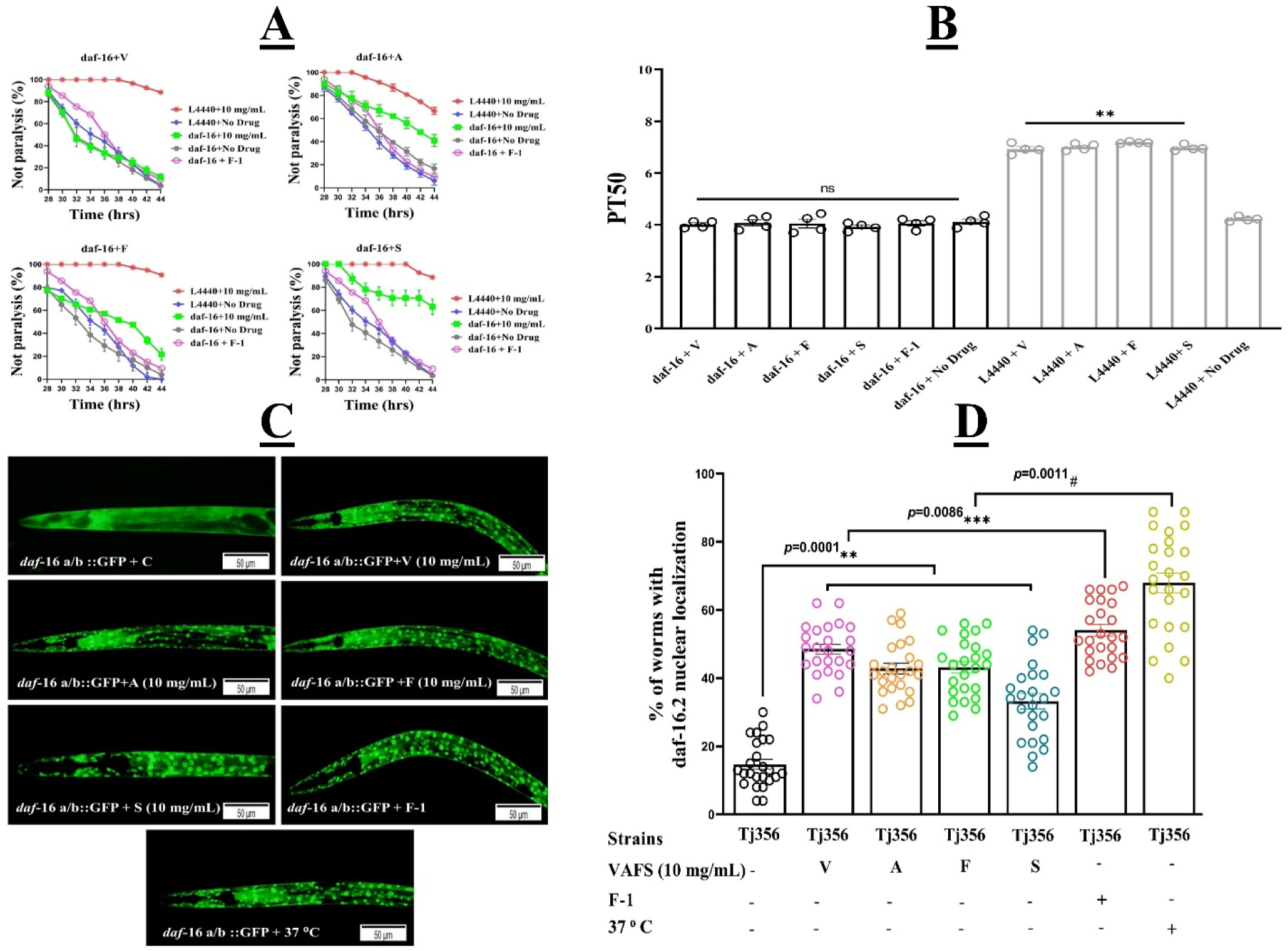
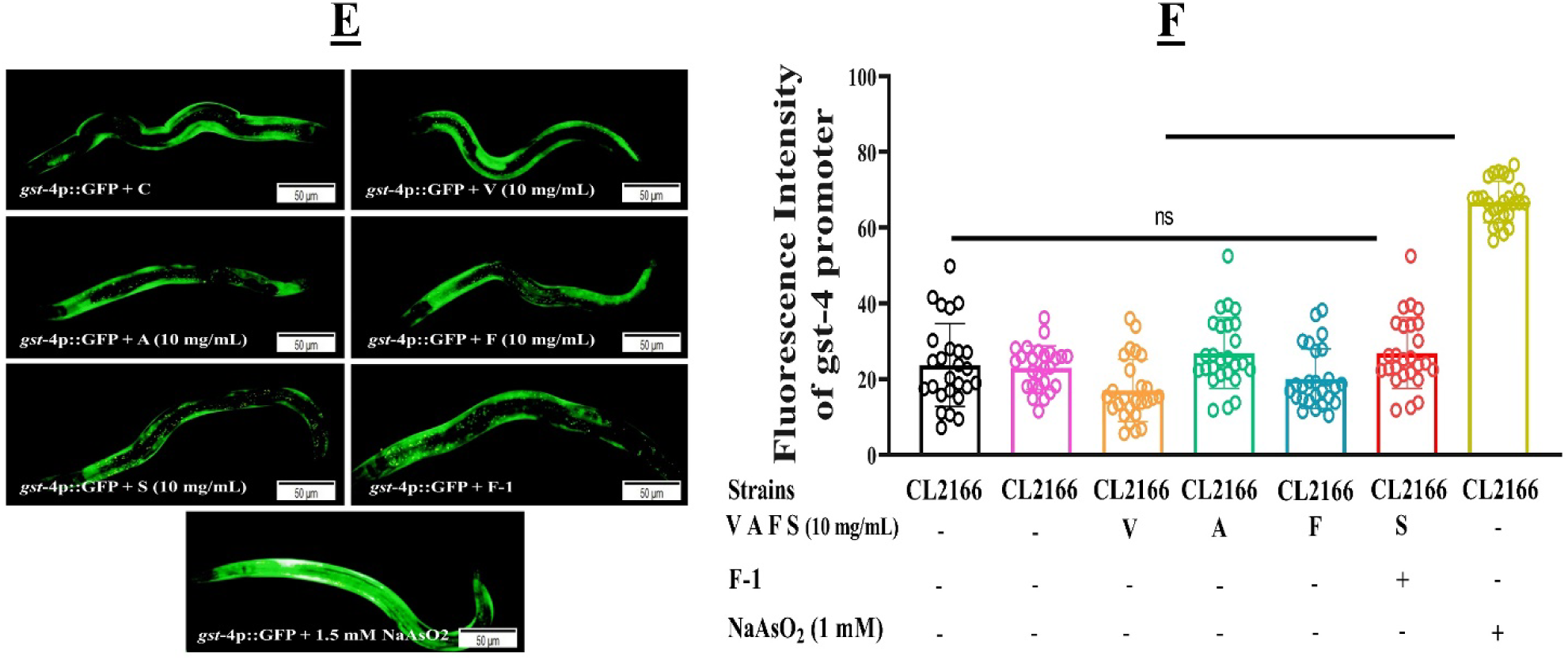
**(A&B).** V, A, F, S on the treatment inhibiting worm AD-like symptom was not via SKN-1 signalling pathway. **(A)** Fractions of not paralysis curves in CL4176(myo-3p::Aβ_1-42_::3′UTR (long) worms after treatment with or without SKN-1 RNAi; (B) PT50s of each treated group in part A, there is a significant difference among these groups when letters are different **(*p*≤0.005). While n represents, no significant difference was found between SKN-1 RNAi/V, A, F, S treated and untreated groups. **(C&D)** Transgenic worms LG333 (*skn*-1b::GFP) were treated with V, A, F, S, and F-1 to observe the *SKN*-1 nuclear localizations. (C) Describing the fluorescent expressions of LG333 after treatment. 1.5 mM sodium arsenide was used as the positive control. The scale bar is 50 μm. (D) Representing the quantitative analysis of fluorescence intensity of treated and untreated worms. Results proved that V, A, F, S, and F-1 did not induce nuclear localization in the treated worms, except the positive control-treated group. n represents no significant difference found between the V, A, F, S, and untreated group, # representing the significant difference between the positive control and untreated group. **(E&F)** Transgenic CL2166 (*gst*-4p::GFP) worms were treated with V, A, F, S, and F-1 to examine the expression of the *gst*-4 gene. (E) Showing the fluorescence expression of CL2166 after V, A, F, S at 10 mg/mL and F-1 treatment. The scale bar is μm. NaAsO_2_ is used here as a positive control. (F) Graphical presentation of V, A, F, S, and F-1 treated and untreated groups. The scale bar is 50. n shows no significant difference between V, A, F, S, and F-1 and untreated groups and failed to enhance the *gst*-4::GFP expressions. # representing the significant difference between positive control (NaAsO_2_) and untreated group.

To verify further whether V, A, F, S, and F-1 could activate the SKN-1 pathway or not, we examined the nuclear localization of intestinal skn-1 in the transgenic worms LG333 (*skn*-1b::GFP). Therefore, we have treated the V, A, F, S, and F-1 with the transgenic worms LG333, which exhibit the intestinal skn-1b activations by the fused GFP reporter translocated from the cytoplasm to the nucleus in response to stress. Our result showed that V, A, F, S, and F-1 did not increase skn-1 nuclear distributions significantly **Fig. 7 (C&D)**.

Glutathione S-transferase (*gst*-4) is a phase II enzyme responding to oxidative stress. At the same time, SKN-1 activation and nuclear translocations play a critical role in regulating downstream stress-responsive gene expressions. To examine whether the expression of *gst*-4 could be induced after V, A, F, S, and F-1 treatment. We selected transgenic worm CL2166 (*gst*-4p::GFP) carrying GFP fused to gst-4 promoter. Later we treated the worms CL2166 with V, A, F, S, and F-1 and observed the results. The result showed that V, A, F, S, and F-1 did not significantly enhance the *gst*-4::GFP expression. Therefore, we concluded that V, A, F, S, and F-1 follow the DAF-16 pathway inhibiting the aberrant Aβ accumulated toxicity but not via SKN-1 in AD models **Fig. 7 (E&F)**.

### 3.7 Zhi-Shi-Wu-Huang tang enhanced the chymotrypsin-like proteasome activity in *C. elegans*

Many factors contributing to PD’s etiologic comprise neuro-inflammation, cellular apoptosis, autophagy deficits, and proteasome dysfunctions. The chymotrypsin-like proteasome system is an essential feature of the proteostasis network. Therefore, to verify whether chymotrypsin-like proteasome activities increase on *Aβ* reductions, we examined the V, A, F, S, and F-1 properties in transgenic CL4176 *C. elegans* and performed a proteasome assay with the fluorogenic substrate (LLVY-R110-AMC). Our assessment showed that drug groups V, A, F, S at 10 mg/mL and F-1 augmented the proteasome’s activities in transgenic CL4176 worms. F-1 showed 18.66 % better proteasomes activities compared with the average of V, A, F, S drug groups. While the level of proteasome activity in CL4176 *C. elegans* is about 48 % lesser than wild-type N2 *C. elegans*. Wild-type N2 is used here as a negative control group. These results verified that drug groups A V, A, F, S, and F-1 augmented the proteasome activity on treatment **Fig. 8 (A)**.

**Figure (8):**
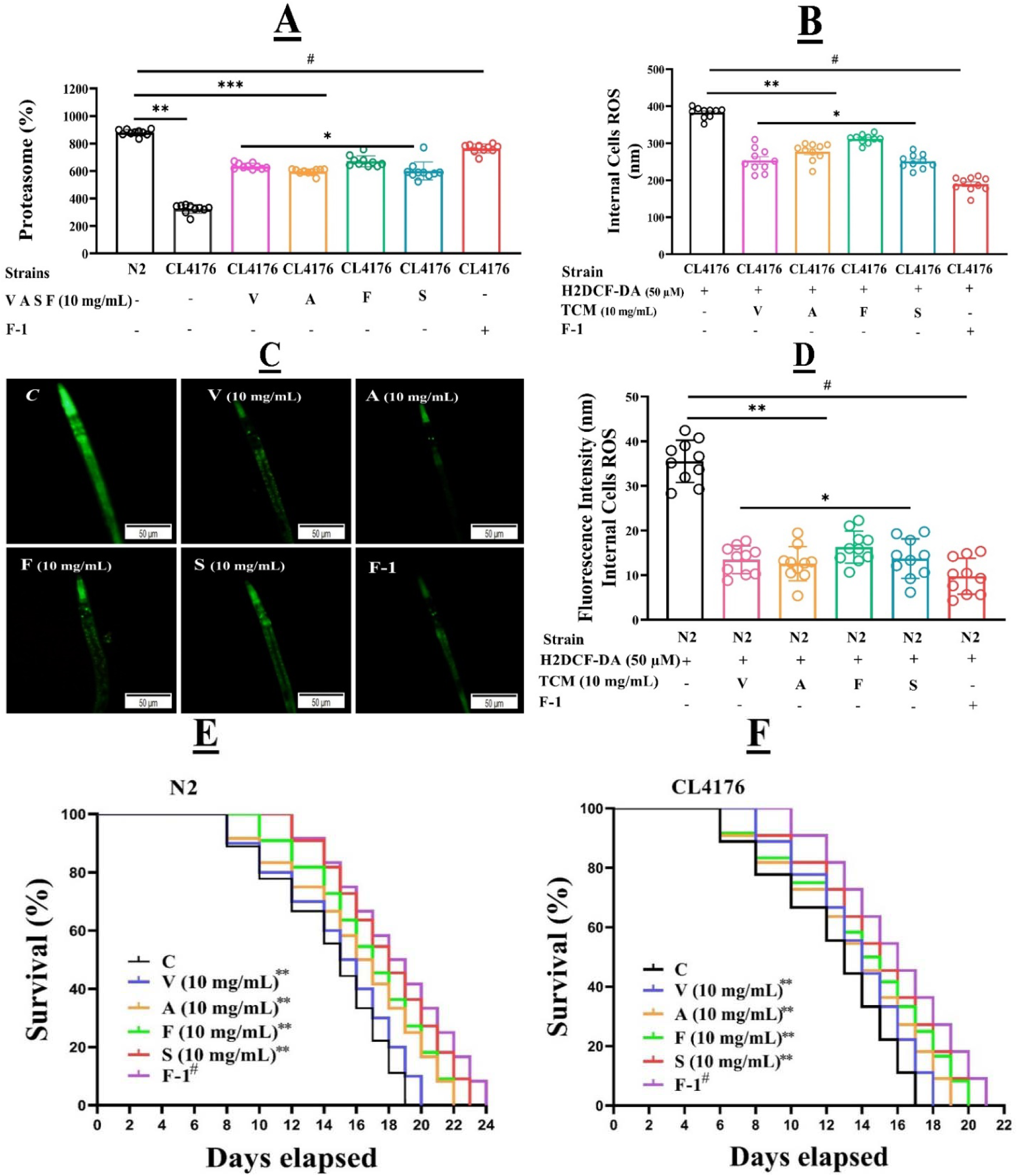
**(A)** TCM drugs V, A, F, S, and F-1 raised the chymotrypsin-like proteasome activities on treatment in CL4176 worms. Proteasome activities were examined after 34 hrs of temperature upshift with or without V, A, F, S at 10 mg/mL dilution. Results show that drug groups V, A, F, S on treatment significantly augmented the proteasome expressions compared to control groups. The data was calculated by mean ± SD (n = 3). ** shows the significant difference of (*p*≤0.005) between V, A, F, S at 10 mg/mL and untreated groups, and # indicating the significant difference between F-1 and control groups. F-1 significantly enhanced proteasome activities much better than single drug treatment (V, A, F, S). It showed that together these four drugs showed decisive neuroprotective actions than a single drug on treatment. **(B)** Explaining the antioxidative role of V, A, F, S, and F-1 on CL4176 worms. Worms were treated with V, A, F, S at 10 mg/mL and F-1. Later supernatant comprises protein and proteasome, transferred to the 96 wells plates after temperature upshift and worm’s lysis. The plates contained fluorogenic substrate (20 µM) and were incubated for 90 min before readings. Data were calculated by mean±SD (n=3). ** shows the significant difference between V, A, F, S treated and untreated groups (*p*≤0.005), # indicating the significant difference between F-1 and control groups. **(C&D)** Describing the antioxidative role of V, A, F, S, and F-1. For that, wild-type N2 *C. elegans* were treated with V, A, F, S at 10 mg/mL and F-1 for 72 hrs. Then worms were treated in 96 wells microtiter plates with fluorogenic substrate H_2_DCF-DA at 50 µM, and fluorescence was observed. Results showed that V, A, F, S, and F-1 significantly reduced the ROS production in worms compared to untreated groups. Data representing the mean±SD (n=15). ** depicting the significance difference of (*p*≤0.005) between V, A, F, S treated groups and the control group. Similarly, F-1 reduced the ROS generation up to 43.34% better than the control group (*p*≤0.005, ***). **(E&F)** Representing the lifespan analysis of wild-type N2 and transgenic CL4176 worms after being treated with TCM (V, A, F, S) at 10 mg/mL. The survival curve was calculated once in two days until all dead, and the cumulative survival curve of three independent tests was calculated via Kaplan-Meier (KM) curve. Control was treated without drugs. Each V, A, F, S drug remarkably enhanced the worms’ lifespans up to 3.66 days compared to the untreated groups (*p*≤0.005, **). At the same time, the F-1 was treated the same way and improved the worm’s lifespan up to 2.33 days better than the average of every single drug on treatment compared with untreated groups (*p*≤0.005, #).

### 3.8 Zhi-Shi-Wu-Huang tang drug reduced ROS in transgenic *C. elegans* on treatment

Previously, the study confirmed that mitochondrial ROS plays a perilous role in inductions of Aβ aggregations in AD (39). Therefore, we have observed the antioxidative properties of V, A, F, S, and F-1 against mitochondrial ROS generations in *C. elegans* models. We confirmed that V, A, F, S, and F-1 drug groups decrease ROS levels in wild-type N2 and transgenic CL4176 *C. elegans* due to their antioxidative activities. We have examined the ROS generations in the treated worms in two ways to confirm the antioxidant activities of drugs. Firstly, in transgenic CL4176 worms, we observed the ROS via spectrophotometer device after incubating the V, A, F, S, and F-1 treated worms in fresh PBS together with H_2_DCF-DA’s microtiter 96 wells plates **Fig. 8 (B).** The results were measured abruptly after H_2_DCF-DA transferring to the experimental groups, and reading was taken every 15 min for an hr. V, A, F, S drugs on treatment reduced the ROS productions compared to the untreated groups (*p* ≤0.005). Secondly, we observed the fluorogenic expression in the fluorescence microscope (BX53) after H_2_DCF-DA incubations of wild-type N2 **Fig. 8 (C&D)**. In the ROS measuring experimental study V, A, F, S, and F-1 significantly reduced the ROS generations in both the observations (*p*≤0.005). Similarly, the F-1 formula drug reduced the ROS productions in both the worms (wild-type N2 and transgenic CL4176) up to 17.44 % on average, much better than untreated groups (*p*≤0.005) **Fig. 8 (B, C&D).**

### 3.9 Zhi-Shi-Wu-Huang tang enhanced the Lifespan of *C. elegans*

Aging is the main factor in developing neurodegenerative disorders and shortening life expectancy (Hung et al., 2010). Therefore, we investigated the effect of V, A, F, S, and F-1 on the lifespan of wild-type N2 and transgenic CL4176 *C. elegans*. Typically, wild-type N2 and transgenic CL4176 *C. elegans* showed a lifespan of about 18.08 ± 0.71 d (*p*≤0.005).

Therefore, at 10 mg/mL, V, F, and S drug groups enhanced the lifespan significantly up to 3.66 days **Fig. 8 (E&F)**. While the F-1 was treated the same way and promoted the life expectancies of worms up to 5.38 days compared to untreated groups. Therefore, our experimental examinations verified that the F-1 formula drug stopped aging in treated worms and enhanced the worm’s life durations up to 2.33 days better than the average of single-drug treatment.

## 4 Discussion

Although there is an ongoing discussion about the Aβ hypothesis, its accumulated toxicity evidence proves the idea that a difference between the production and clearance of Aβ is one of the critical starting events in AD progressions. Aβ oligomers elevations have been found to drive the growth of neuroinflammation, synaptic loss, neuron cell death, and oxidative stress. Therefore, there is considerable interest in finding plant-based anti-Aβ drugs to attenuate Aβ-induced toxicity. TCM has been used for thousands of years with high efficacy and low toxicity metabolic records (40). The TCM-derived therapeutics, including herbal decoctions and individual active constituents, have been used in clinics for a long time (8). To prevent and treat AD progressions, we preferentially selected the top four HSP70 promoter activator TCMs from the 35 traditional prescriptions, comprising Valeriana jatamansi Jones (Zhi Zhu Xiang, “V”), Acorus tatarinowii (Shi Chang Pu, “A”), Fructus Schisandra Chinensis (Wu Wei Zi, “F”), and Scutellaria baicalensis (HuangQin, “S”). These four V, A, F, S proved the benefit of curing via heat shock protein (HSP) promoter activations previously done in our laboratory. Heat shock proteins can prevent false folding and protein aggregation re-fold the misfolded protein, especially the heat shock protein HSP70 (41). Studies have shown that HSP70 plays a vital role in reducing protein accumulations of α-Syn, Aβ_1-42_, restoring tau protein in vivo balance, reducing oxidative stress, and inhibiting nerve inflammations (10, 42). Therefore, in our present work, we confirmed that V, A, F, S and the newly discovered formula named “F-1” significantly inhibited Aβ-induced paralysis and displayed neuroprotective effects in transgenic CL4176 strains. This suggests that V, A, F, S at 2.5 mg/mL, 5.0 mg/mL, 10 mg/mL and F-1 could potentially treat AD. To begin with, we identify the V, A, F, S, and F-1 toxicity effects on *C. elegans* physiology. OP50 bacterial strain (E. coli) was used as a food source for *C. elegans* (43). These *C. elegans* based assays could be influential for a quick assessment, cost-effective, and screening larger numbers of novel neuroprotective drugs in a short period (44). Our food clearance results confirmed that V, A, F, S at 2.5 mg/mL, 5.0 mg/mL, 10 mg/mL, and F-1 drug groups in s-medium did not affect the *C. elegans* growth considered to be non-toxic than F, S drug groups at 20 mg/mL, and 30 mg/mL concentrations groups **Fig 1**. Aβ_1-42_ aggregative toxicity is one of the pathological hallmarks of AD. To prove the anti-Aβ_1-42_ effects of V, A, F, S, and F-1, we have selected transgenic CL4176 worms that showed paralysis phenotypic expressions on temperature upshift to 25 □C. Results verified that V, A, F, S, and F-1 improved Aβ-induced paralysis in the transgenic CL4176. In addition, to confirm whether V, A, F, S, and F-1 effects non-specifically on Aβ expressions or exclusively act on Aβ aggregative toxicity, CL2179 worms were used as a transgenic background control (45). Expectedly, the GFP fluorescence intensity of treated V, A, F, S, and F-1 groups revealed no significant difference with untreated groups **Fig 2 (B&C).** Results verified that V, A, F, S, and F-1 significantly alleviated exogenous 5-HT hypersensitivity tempted by Aβ accumulated toxicity **Fig. (3)**. It is well known that Aβ depositions (monomers, oligomers) prompt neuron death and are responsible for AD-related amnesia (46). Likewise, Aβ deposits are closely related to the severity of memory impairment in AD patients (47). The amyloid hypothesis postulates that Aβ depositions encourage the course of AD pathogenicity (48). Therefore, decreasing the Aβ deposits is one of the most promising therapeutic approaches to treat AD (49). Our results supported V, A, F, S, and F-1 strongly ameliorating Aβ deposits in the transgenic CL2006 (Aβ_1-42_) worms **Fig. 4 (A&B)**. After checking the V, A, F, S, and F-1 effects against Aβ oligomeric expressions, we verified the anti-Aβ accumulative effects by immunoblotting assay **Fig 4 (C&D)** V, A, F, S and F-1 could eliminate the accumulated Aβ_1-42_ toxicity by activating the DAF-2/16 pathway in transgenic worms CL4176 verified via RNA interference (RNAi) assay (50). For that, we have knocked down the expression of the transcription factor DAF-16 by RNAi in transgenic worms CL4176 and experienced the inhibitory effect of V, A, F, S on worms paralysis **Fig. 5 (A&B).** Similarly, DAF-2/16 pathway activations in Tj356 worms were confirmed on V, A, F, S treatment. DAF-16 is a transcription factor and a primary component in the DAF-2/DAF-16 Insulin-like Signaling Pathway(35, 51). The activation of DAF-16::GFP gave higher fluorescence expressions of genes in responses to a wide range of stressors. Our study showed that V, A, F, S, and F-1 significantly activated DAF-16::GFP, translocate from the cytosol to nucleus expressions in Tj356. Cytosolic to nuclear translocation confirmed that DAF-16 is required to delay the paralysis effect of V, A, F, S **Fig. 5 (C&D).** HSPs are Heat Shock Proteins with 12-43 kDa molecular weight (52). In the case of thermal and oxidative stresses, HSP proteins respond to neutralize anxiety (53). However, HSPs inhibit the accumulations of numerous toxic proteins, such as ploy Q and Aβ (54). HSP16.2 can suppress Aβ toxicity by assisting abnormal protein refolding and degradation in AD transgenic Tj375 worms. V, A, F, S, and F-1 on treatment did not increase the expression of hsp-16.2, representing that Aβ oligomeric toxicity reduction was not interceded by molecular chaperon HSP-16.2. However, V, A, F, S decreased the hsp-16.2 up-regulation induced by Juglone. Since Juglone is a stress-inducer (55), it proved that V, A, F, S, and F-1 could reduce Aβ toxicity through their antioxidant properties **Fig. 6 (A&B).** Superoxide dismutase (SOD-3) protects transgenic worms CF1553 from oxidative stress and suppresses the Aβ toxicity. In *C. elegans*, *SOD*-3 is located downstream of DAF-16 (56). The recent study showed that V, A, F, S, and F-1 enhanced the *SOD*-3 expressions **Fig. 6 (C&D)**. SKN-1 pathway is a transcriptional factor parallel to the DAF-16 pathway, can also promote the expression of stress-responsive genes to prevent the Aβ_1-42_ toxicity (30). In our experiments of SKN-1 RNAi genes, knockout V, A, F, S, and F-1 did not affect delaying paralysis of anti-AD actions. While SKN-1 did not express nuclear translocations after V, A, F, S, and F-1 treatment in transgenic LG333 (skn-1b::GFP), and CL2166 (gst-4p::GFP) worms **Fig. 7** (47). The chymotrypsin-like proteasome system (CPS) enhanced cell survival on damage under environmental stress conditions (57). The 26S proteasome complex of *C. elegans* includes the 20S catalytic centre (58). This massive protein complex possesses 19S regulatory caps and active proteolytic centres to identify protein substrate poly-ubiquitinated overwhelmed for deficiencies (59). The effect of V, A, F, S, and F-1 is associated with higher expressions of proteasome activity at the cellular level. Ectopic expression of proteasome units via chymotrypsin-like activities is sufficient to confer proteotoxic stress resistance and extend lifespan (33). However, our chymotrypsin-like activity assay demonstrated that V, A, F, S, and F-1 actively enhanced the proteasome 20S functions on treatment **Fig 8 (A)**. Therefore, improvement in Aβ-induced pathological symptoms by V, A, F, S, and F-1 treatment could be due to the proteasomes pathway, but further research needs to be done in the future. Indeed, toxic Aβ is closely related to the CPS. On the one hand, impairments of the CPS lead to low abnormal Aβ clearance and subsequently accelerate the aggregations of Aβ in AD (60).

As we know, Aβ toxicity generates ROS at the mitochondrial level and could cause AD-like symptoms (61). AD is susceptible to oxidative stress via ROS (62). At the same time, ROS is the main contributor to apoptosis and hyperphosphorylation (63). Therefore, we treated the wild-type N2 and transgenic CL4176 worms with V, A, F, S, and F-1 to confirm their antioxidative properties. Our assessment showed that V, A, F, S, and F-1 significantly declined the live animals’ cellular and cellular ROS productions on treatment **Fig 8 (B, C&D)**. AD is an age-related disorder (64). Therefore, we examined the V, A, F, S, and F-1 effects on *C. elegans* lifespan. The purpose was to identify the V, A, F, S, and F-1 anti-aging properties. So, on treatment, V, A, F, S, and F-1 significantly prolonged the worms’ lifespan up to 4-5 days (**Fig. 8 (E&F)**. The treatment of V, A, F, S, and F-1 increased organism stress resistance to secure against oxidative stress prompted by abnormal Aβ proteins. Moreover, Aβ (oligomer) accumulated toxicity is closely related to the severity of memory impairments in AD patients. Our results supported that V, A, F, S, and F-1 strongly ameliorating Aβ toxicity in the transgenic worms integrated with human Aβ_1-42_.

## 5 Conclusion

Zhi-Shi-Wu-Huang Tang, which comprises Valeriana jatamansi (V), Acori talarinowii (A), Scutellaria baicalensis (S) Fructus Schisandrae (F) is a newly discovered named “F-1” with multiple therapeutic actions. These four TCM are the best HSP 70 promoter activator and express heat shock proteins (HSP-70). HSP-70 can prevent false folding and protein aggregations. F-1 was discovered and calculated based on V, A, F, S anti-Aβ accumulative effects in AD transgenic worms via orthogonal paralysis tests. Our experimental assessment confirmed every drug V, A, F, S at various dilutions, and F-1 reduced Aβ aggregative toxicity by activating the Insulin Daf-16 signaling pathway. We also examined that V, A, F, S, and F-1 reduced the ROS generations and augmented the chymotrypsin-like proteasome expressions in transgenic animals. Therefore, we predict that V, A, F, S, and F-1 could activate the oxidative pathway or CPS pathway other than DAF-16 in treated animals to show their anti-AD actions. But, this study will be done in our future projects.

## Author’s Declarations

### Ethical statement

n/a

### Consent for publication

n/a

### Funding

1- Natural Science Foundation of Gansu Province of China, 20JR10RA596,20JR10RA756

2- Talent innovation and entrepreneurship project of Lanzhou City, 2020-RC-43.

### Author’s contributions

All authors contributed to the concept of this study. All authors provided critical feedback and helped shape the research, analysis, and manuscript.

### Availability of data and materials

The aggregate data supporting findings contained within this manuscript will be shared upon request and submitted to the corresponding author.

