## Supplementary data for "Zhi-Shi-Wu-Huang attenuates Aβ toxicity in *Caenorhabditis elegans* Alzheimer’s disease models via modulating insulin DAF-16 signaling pathway"

**Supplementary experimental data**

### 1 HPLC Analysis of V, A, S, F

#### 1.1 **HPLC required Reagents.**

Acetonitrile 80 % (HPLC grade) and Ultrapure water (20 %) were obtained using the Milli-Q system (Millipore, Bedford, MA, USA) was used in the experiments. Methanol (HPLC grade) was used to clean the column. Eight standards for quantitative analysis were purchased from the National Institution for Food and Drug Control.

#### 1.2 **Analytical Conditions and Instrumentation.**

HPLC system 1200 series (Agilent Technologies, USA) equipped with Chemstation B.03.02 software (Agilent Technologies, USA) comprised a quaternary solvent delivery pump, an online vacuum degasser, an autosampler, a thermostatic compartment, and a UV detector were used for chromatographic analysis. All separation processes were performed using a C_18_ column (5.0 m particle size with 250 mm ×4.6 mm i.d) Kromasil.

#### 1.3 **Mobile phase A was acetonitrile, and phase B was water.**

The linear gradient condition (30 % B for 0 min to 8 min; 30% to 20% B for 8 min to 12 min; 20% to 10% B for 12 min to 20 min; 10% B for 20 min to 30 min) was applied for the separation process. The eight selected standards can be analysed completely by using this procedure. The flow rate was 0.8 mL/min, and the column temperature was 25 ^∘^C, which was maintained for the entire experiment. The eluate was monitored at 314 nm, and the injection volume was 30 µL. The peak identification was based on the retention time and UV spectrum against the standard presented in the chromatogram.

#### 1.4 **Preparation of the standard solution.**

Quantification was based on the standard external method. The stock solutions of each standard were prepared by dissolving in methanol. The solutions were separately and precisely prepared as follows: Caffic acid (0.0025 g), Valtrate (0.0025 g), β-asarone (0.0025 g), α-asarone (0.0025 g), Gallic acid (0.0025 g), Schizandrol A (0.0035 g), Baicalin (0.0025 g), and Chrysin (0.0025 g) were placed in centrifugal tubes, in which 2, 2, 2, 2, and 2.5 mL of methanol were then added, respectively. After 10 min ultra-sonication, filter with a syringe filter before analyzing the HPLC. All of the solutions were stored at −4^∘^C.

#### 1.5 **Working standard solutions preparations.**

The working standard solutions were prepared using the stock solutions. The appropriate stock solutions and methanol were mixed well. Finally, the standard working solutions (800, 261.5, 255.7, 200, 200 g/mL, 200 g/ml, 150 g/ml) of Caffic acid, Valtrate, β-asarone, α-asarone, Gallic acid, Schizandrol A, Baicalin, and Chrysin were prepared. Subsequently, the mixed standard work solutions (1.25 g/mL to 300 g/mL) of Caffic acid, Valtrate, β-asarone, α-asarone, Gallic acid, Schizandrol A, Baicalin, and Chrysin were prepared by using the working standard solutions. All of the solutions were stored at −4 ^∘^C.

#### 1.6 **V, A, S, F sample preparations for HPLC.**

The powdered samples of each drug were refluxed using ultra water for 2 hrs; this process was repeated at 90 °C. The extracts were combined before being filtered and then diluted with water until the final volume of 0.25 mg herbs/mL was reached. The diluted solution was filtered through a syringe filter (0.25 m) and stored at −4 °C before injection.

#### 1.7 **Zhi-Shi-Wu-Huang drug preparation**

Different combinations of V, A, S, and F for orthogonal experiments were prepared according to **Table 2**. Then we treated transgenic *C elegans* (OW13, CF1553) with **Table 3** dilations. After treatment and analysis, the orthogonal experimental dilutions help prepare the formula drug at various V, A, F, S and F ratios. The results of orthogonal experiments and the procedure for calculating the formula from V, A, F, S are given in the supplementary file (Supplementary data Fig: S01). We successfully computed the formula and called in Chinese Zhi-Shi-Huang-Wu. The Zhi-Shi-Wu-Huang in 8:4:2:1 ratio (specifically at V: 20 mg/mL, A: 10 mg/mL, : 5 mg/mL, F: 2.5 mg/mL) named as **F-1**.

### 2 V, A, F, S, and F-1 HPLC experimental outcomes

#### 2.1 **Wavelength optimization.**

The eight mentioned standards were scanned using the UV detector between 190 and 400 nm. The maximum absorbance values of Caffic acid, Valtrate, β-asarone, α-asarone, Gallic acid, Schizandrol A, Baicalin, and Chrysin were 314, 310, 270, 280, 330 nm, 254, 280, and 370, respectively. Given the serious end absorption near 270 nm, the detection wavelength was set at 270 nm, in which all of the compounds have an apparent absorption band. The results are shown in **Figure 1.**

#### 2.2 **Detector optimization.**

The UV detector can detect only one wavelength at a time. Therefore, measuring the standard's characteristic absorption and retention time was easier. A general assumption can be made using the characteristic absorption and retention time. In this way, the error rate is unusually reduced and suitably used to valuation each chromatographic peak of the corresponding samples.

#### 2.3 **Mobile phase optimization.**

Several organic solvents in different ratios resulted in different peak separations. The composition and ratios of the mobile phase were optimized using different organic solvents, including acetonitrile and water in different concentrations. Methanol was used to clean the column. In the present study, the mobile phase composition was selected as the mobile phase for the simultaneous analysis because of better resolution and shorter analysis time than other phases. Typical HPLC-PDA chromatograms of the standard samples are presented in (**Supplementary figure 1)**.

#### 2.4 **Stability.**

The mixed working standard solutions of different concentrations were analyzed at 0, 3, 6, 9, 15, 21, and 24 h at room temperature through the method's stability. The peak area was recorded, and RSD was calculated. The RSDs were 0.30%, 0.31%, 0.30%, 0.37%, and 0.48%, 0.54 %, 0.67 % and 0.71 % (n= 8), which indicated that the established method was stable in a day.

#### 2.5 **Repeatability.**

Two portions were precisely weighed from each V, A, S, F and formula drug sample; the Caffic acid, Valtrate, β-asarone, α-asarone, Gallic acid, Schizandrol A, Baicalin, and Chrysin contents were detected through the developed method; the RSDs of the samples were 2.80 %, 7.284 %, 3.39 %, 3.29 %, 2.8 4%, 3.19 %, 2.81 % and 3.22 % respectively. Standard contents were .246 mg/mL, .280 mg/mL, .270 mg/mL, .253 mg/mL, .293 mg/mL, .262 mg/mL, and 0.26 mg/.253 mg/mL, respectively.

#### 2.6 **Applications.**

Using the new and modern method, the analytic contents of the four selected TCM and formula drugs from various regions of China were measured. Marked differences were observed in the analytic content of TCM samples. This TCM has the highest amount of Valtrate (7.284 %) in V, α-asarone (3.297 %) in A, Schizandrol A (3.19 %) in F, and Baicalin (2.88 %) in S were relatively high. The analytic entities were almost similar to the other area of herbs (**Supplementary figure 2)**.

### 3 V, A, S, F and F-1 enhanced the Chemotaxis Behavior

The chemotaxis response in *C. elegans* is mediated by activating several sensory neurons and interneurons to stimulate the motor neurons (Hobert, 2003). The Chemotaxis Index (CI) measures the fraction of worms that can arrive at the location of the attractants (Matsuura et al., 2005). To investigate the effect of V, A, S, F and F-1 on the performance of chemotaxis behavior, we applied the chemical benzaldehyde as an attractant and ethanol as a control, both containing sodium azide, which paralyzes the worms on contact (supplementary figure 1). The chemotaxis index was scored in the transgenic and control strain on day 4. Supplementary figure 1 shows that the transgenic mutant CL2355 exhibits a significant reduction of CI compared with the control strain CL2122, or no Aβ strain untreated. V, A, S, F feeding of the control strain did not affect their CI, but it significantly normalizes the reduced CI in the neuronal A transgenic strain; n = 4; p 0.01; a total of 40 worms was used in each assay).

#### 3.1 **Results**

##### 3.1.1 **Wavelength Optimization.**

The eight mentioned standards were scanned using the UV detector between 190 and 400 nm. The maximum absorbance values of Caffic acid, Valtrate, β-asarone, α-asarone, Gallic acid, Schizandrol A, Baicalin, and Chrysin were 314, 310, 270, 280, 330 nm, 254, 280, and 370, respectively. Given the serious end absorption near 270 nm, the detection wavelength was set at 270 nm, in which all of the compounds have an apparent absorption band. The results are shown in Figure 1**.**

##### 3.1.2 **Detector Optimization.**

The UV detector can detect only one wavelength at a time. Therefore, measuring the standard's characteristic absorption and retention time was easier. A general assumption can be made using the characteristic absorption and retention time. In this way, the error rate is unusually reduced and suitably used to valuation each chromatographic peak of the corresponding samples.

##### 3.1.3 **Mobile Phase Optimization.**

Several organic solvents in different ratios resulted in different peak separations. The composition and ratios of the mobile phase were optimized using different organic solvents, including acetonitrile and water in different concentrations. Methanol was used to clean the column. In the present study, the mobile phase composition was selected as the mobile phase for the simultaneous analysis because of better resolution and shorter analysis time than other phases. Typical HPLC-PDA chromatograms of the standard samples are presented in (supplementary figure 1).

##### 3.1.4 **Stability.**

The mixed working standard solutions were analyzed at 0, 3, 6, 9, 15, 21, and 24 h at room temperature to evaluate method stability. The peak areas of eight analytes were recorded and the relative standard deviations (RSDs) were calculated. The RSD values ranged from **0.30% to 0.71% (n = 8)**, indicating that the developed method exhibited good stability within 24 h.

| Compound | RSD (%) (n=8) |
| --- | --- |
| Caffeic acid | 0.30 |
| Valtrate | 0.31 |
| β-Asarone | 0.30 |
| α-Asarone | 0.37 |
| Gallic acid | 0.48 |
| Schizandrol A | 0.54 |
| Baicalin | 0.67 |
| Chrysin | 0.71 |

##### 3.1.5 **Repeatability.**

Two portions of each sample (V, A, S, F, and the formula drug) were accurately weighed and analyzed using the developed HPLC method. The contents of caffeic acid, valtrate, β-asarone, α-asarone, gallic acid, schizandrol A, baicalin, and chrysin were determined. The RSD values ranged from 2.80% to 7.28%, demonstrating acceptable repeatability of the method.

| Compound | Standard Concentration (mg/mL) | RSD (%) |
| --- | --- | --- |
| Caffeic acid | 0.246 | 2.80 |
| Valtrate | 0.280 | 7.28 |
| β-Asarone | 0.270 | 3.39 |
| α-Asarone | 0.253 | 3.29 |
| Gallic acid | 0.293 | 2.84 |
| Schizandrol A | 0.262 | 3.19 |
| Baicalin | 0.260 | 2.81 |
| Chrysin | 0.253 | 3.22 |

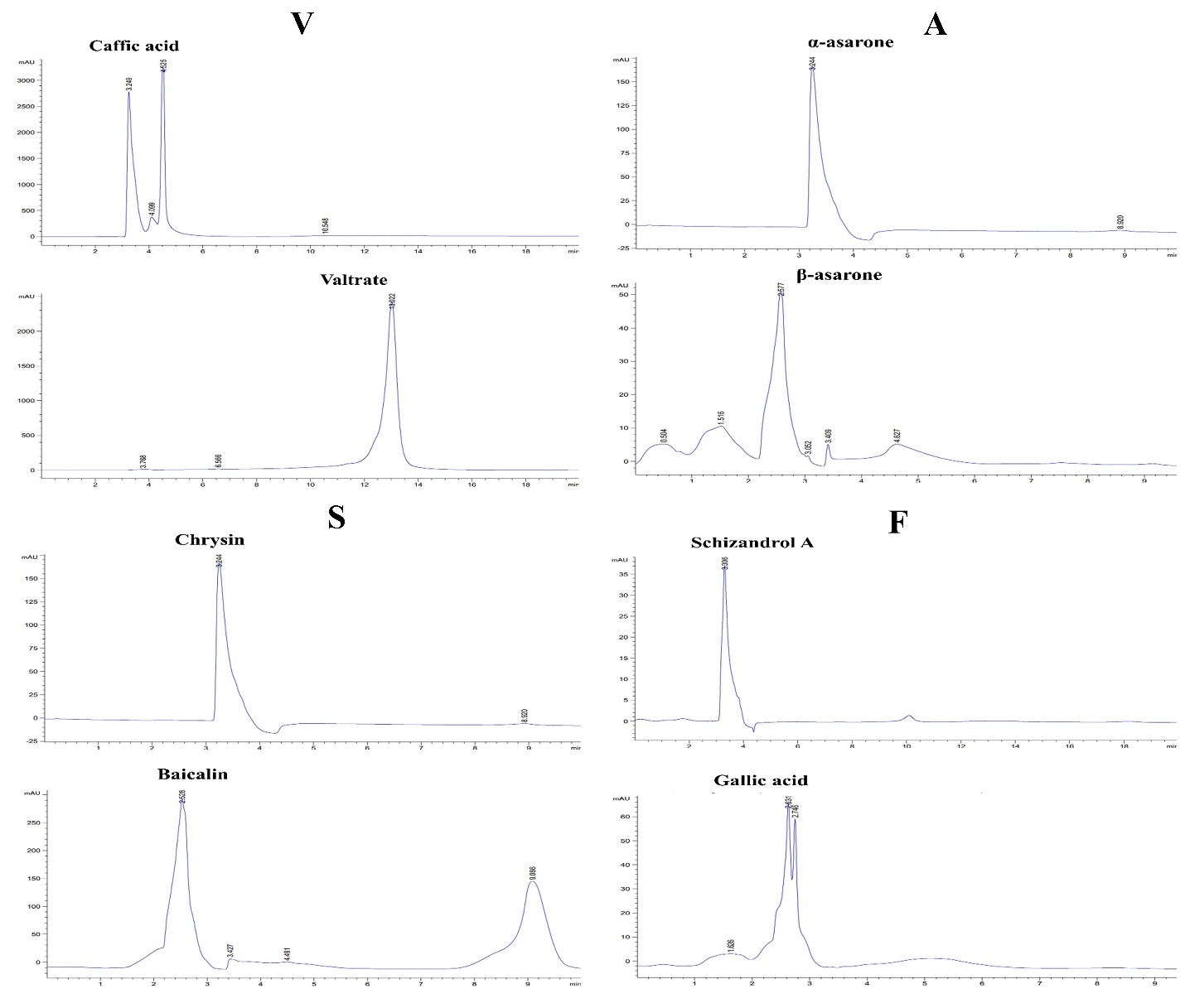
**Supplementary figure 1:** Explaining the chromatograms of particular standards for each TCM Caffic acid, Valtrate, β-asarone, α-asarone, Gallic acid, Schizandrol A, Baicalin, and Chrysin under optimal chromatographic conditions.

##### 3.1.6 **Applications.**

Using the new and modern method, the analytic contents of the four selected TCM and formula drugs from various regions of China were measured. Marked differences were observed in the analytic content of TCM samples. This TCM has the highest amount of Valtrate (7.284 %) in V, α-asarone (3.297 %) in A, Schizandrol A (3.19 %) in F, and Baicalin (2.88 %) in S were relatively high. The analytic entities were almost similar to the other area of herbs (Supplementary figure 2).

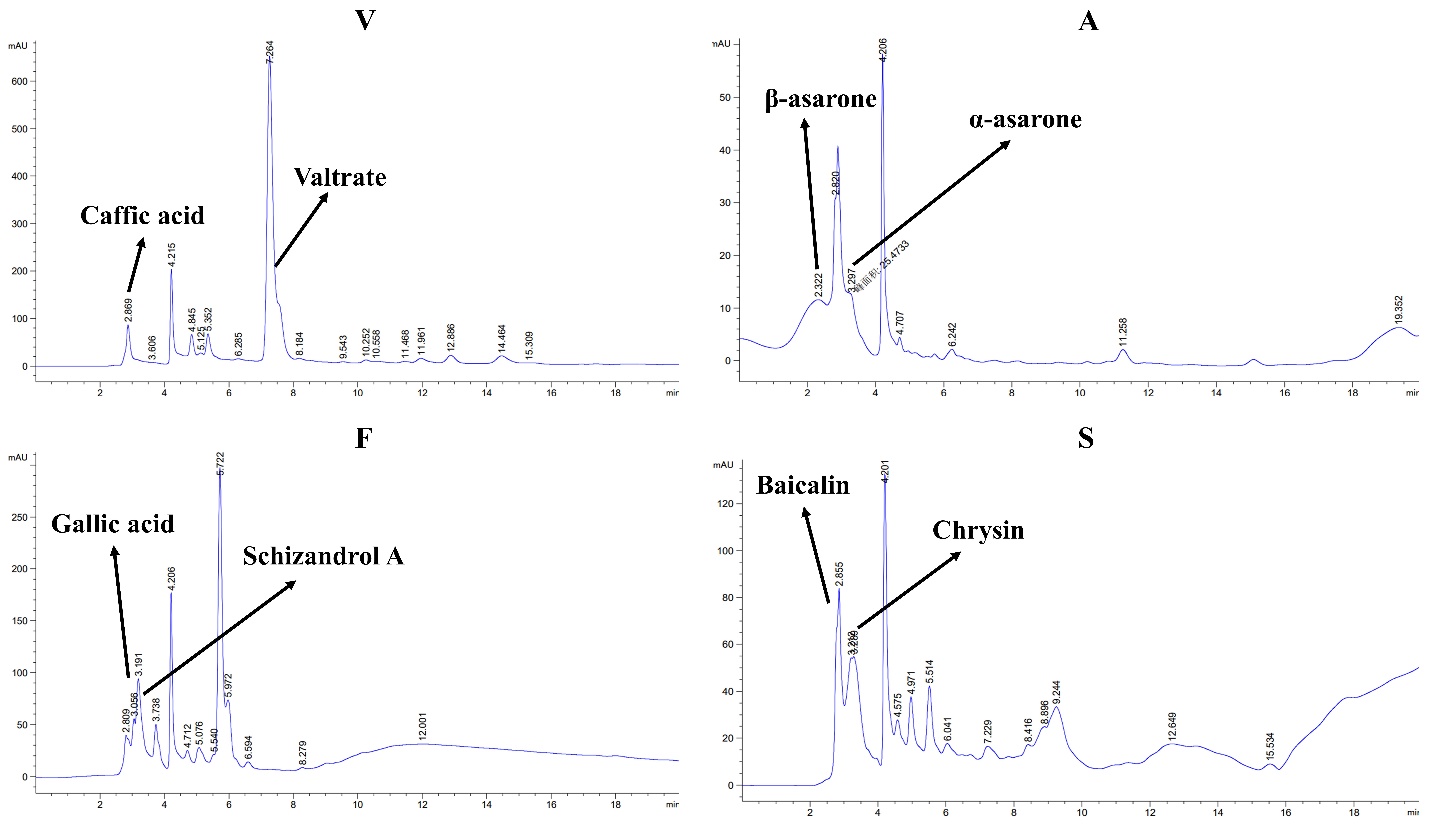
 **Supplementary figure 2:** Representing the chromatograms of V, A, F, S TCM with the percentage of standard-specific standards represented by peak values.

### 4 Orthogonal experimentations

### Percentage of V, A, F, S drug calculate formula drug (Zhi-Shi-Wu-Huang)

We performed an orthogonal experimental study on various Zhi-Shi-Wu-Huang dilutions to identify the anti-Aβ toxicity effects of each drug dilution in transgenic CL4176 worms via paralysis assay. Orthogonal dilutions of each drug (V, A, F, S) are given in **Table: 3**. Similarly, based on orthogonal assessment, Zhi-Shi-Wu-Huang that computed on the ratio comprised of V, A, F, S at an 8:4:2:1 ratio, specifically V: 20 mg/mL, A: 10 mg/mL, F: 5.0 mg/mL, and S: 2.5 mg/mL had shown significant anti-AD effects on transgenic *C. elegans,* named as Formula-1(F-1) in this study. The F-1 was selected based on each V, A, F, S drug neuroprotective effect and toxicity to the treated AD *C. elegans* models. Later, we compared F-1 and every drug’s (V, A, F, S) neuroprotective functions in *C. elegans models* in the following anti-Aβ approaches.

### 5 Zhi-Shi-Wu-Huang’s food clearance assay

To determine the effective and nontoxic drug concentration of V, A, F, and S extract, we conducted a food clearance test in wild-type N2, and transgenic CL4176 (myo-3p::Aβ_1-42_::let-851 3'UTR) worms at 0 mg/mL, 2.5 mg/mL, 5.0 mg/mL, 10, 20 mg/mL, and 30 mg/mL concentrations in S-medium. The purpose was to determine the toxic role of treated drugs on the worm’s physiology and behaviours. We observed worms treated with F and S at 20mg/mL and 30 mg/mL dilution were showed toxic behaviour, leading to the reduced body size with smaller numbers of offspring, even causing the death of adults at day’s interval and lowering the food clearance curve. All other drug extract-treated groups were alive with normal physiological conditions up to 10 mg/mL and did not show a significant difference compared to the control groups (Figure 1). Therefore, V, A, F, S dilutions at 10 mg/mL were considered to treat *C. elegans* for further experimental study. At the same time, the non-toxic concentration results of the F-1 decoctions are given together with V, A, F, S drug dilutions (Supplementary figure 3).

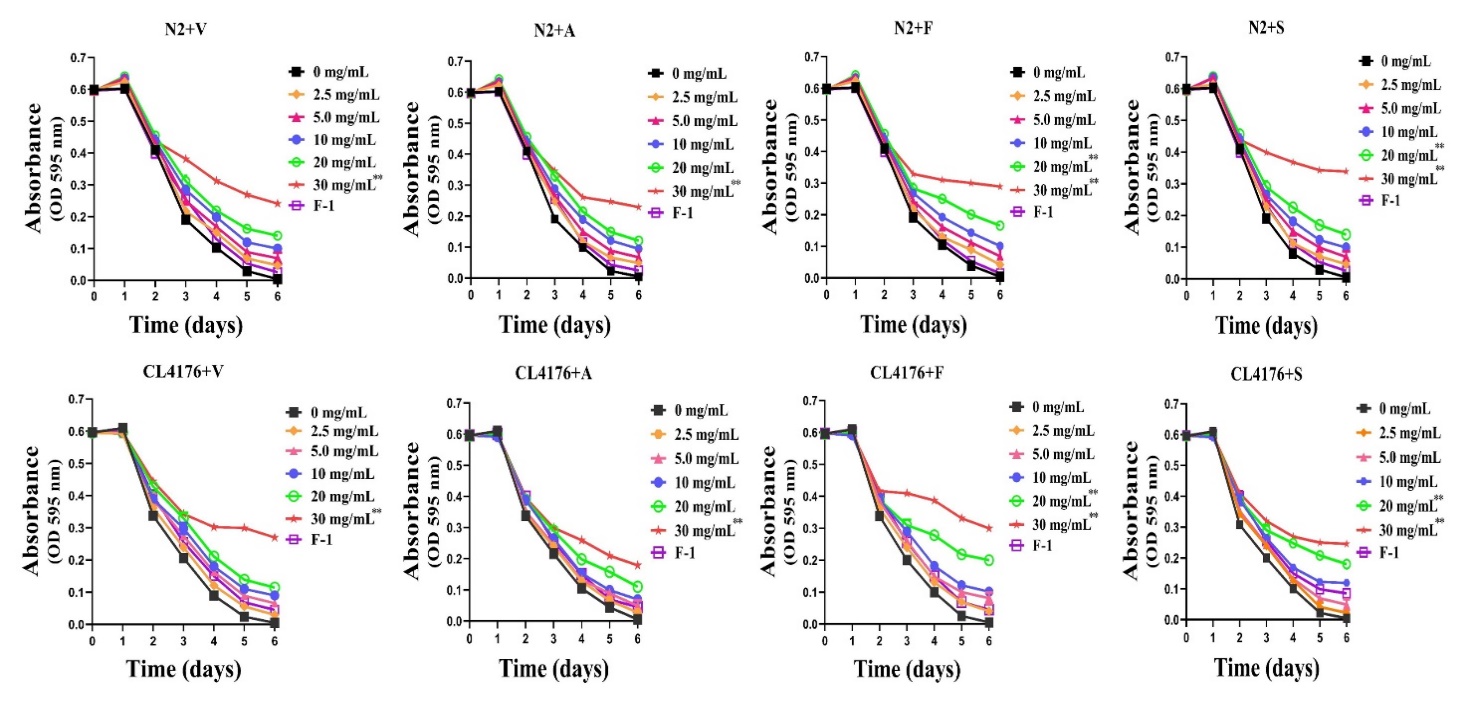
**(Figure 1):** Explaining the results of the food clearance test of V, A, F, S extract in the pharmacological *C. elegans* models of AD. A food clearance test was performed to evaluate the non-toxic concentrations of selected drugs against treated models. Briefly, in a 96-well plate, synchronized L1 worms of wild-type N2, and transgenic CL4176 were cultured with OP50 bacterial strain (OD A_595_ = 0.6 absorbance’s) feeding medium containing drugs V, A, F, S at 2.5 mg/mL, 5.0 mg/mL, 10 mg/mL, 20 mg/mL, and 30 mg/mL concentrations and F-1 for 6 days. The OD value of each treatment was measured and recorded on the day interval. Results showed that F and S at 20 mg/mL and 30 mg/mL showed drug toxicity and caused *C. elegans* to become thin and slender in growth, even leading to death. All the selected V, A, F, S drugs showed non-toxic dilutions up to 10 mg/mL and the F-1 drug group to treat *C. elegans*. While the only significant difference was found between the V, A, F, S dilutions (20 mg/mL, 30 mg/mL) and the control groups in treated models (wild-type N2 and transgenic CL4176) is *p*≤0.005 (**).

### 6 V, A, S, F and F-1 reduced the α-Syn aggregations in transgenic *C. elegans*

To examine the expression and aggregation of α-Syn, we used the *C. elegans* strain OW13, which was constructed with human α-Syn fused with a yellow fluorescent protein (YFP) under control of the unc-54 promoter where gene expression occurred in the body wall of muscle cells (Van Ham et al., 2008). OW13 strain's advantages lead to the high efficiency of expression and the PD-like progressive defects of *C. elegans* motility. Therefore, it demonstrated the in vivo aggregation

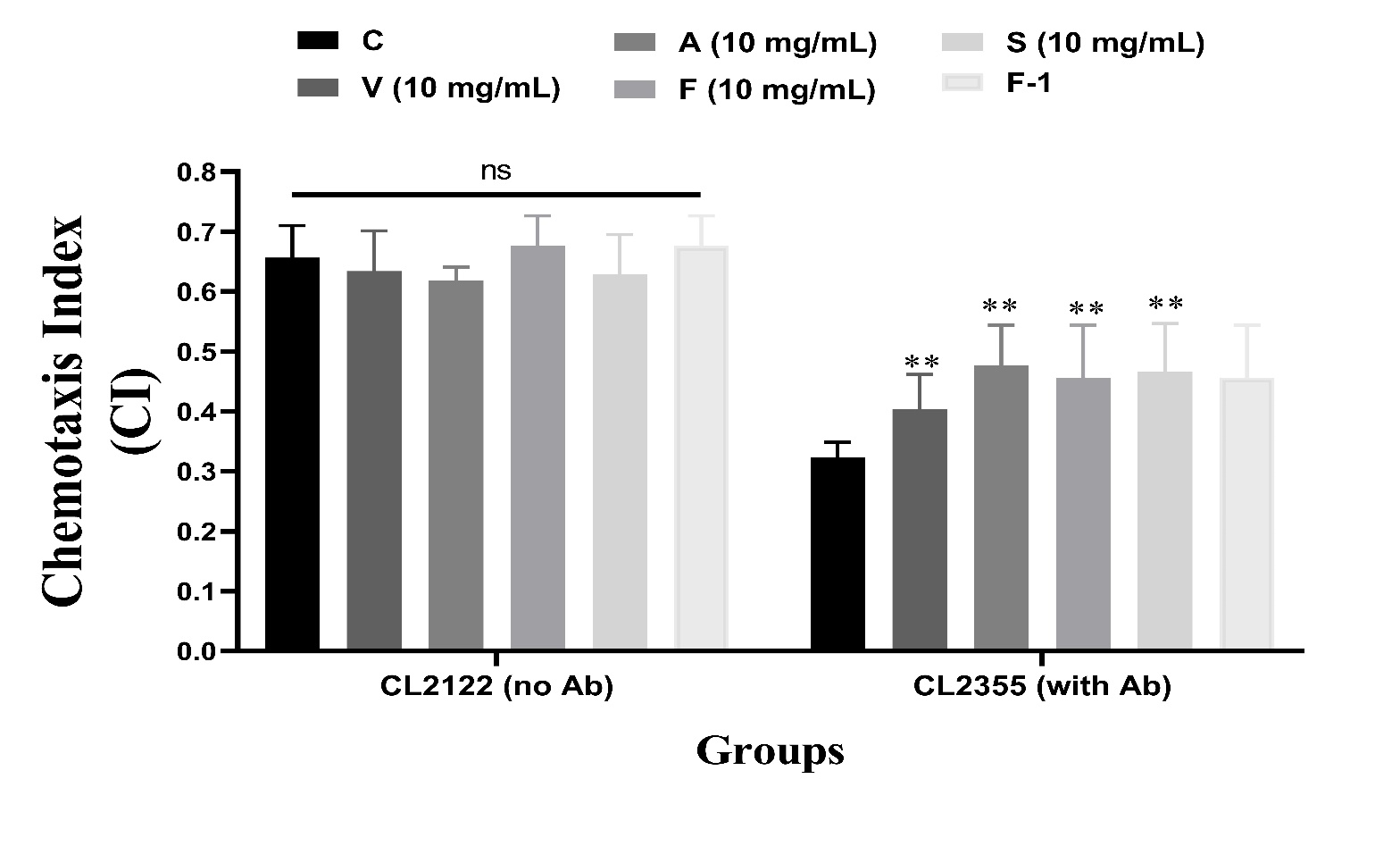
 **Supplementary figure 3:** Depicting a graphical diagram of chemotaxis assay at 10 mg/mL concentration of selected 4 TCM (V, A, S, F) and a formula drug F-1. For methodology, please go to material and methods. Results described a significant decrement of CI between no Aβ + neuronal Aβ containing *C. elegans* with *p*<0.005 and showed a significant difference between drug-treated groups with control groups *p*<0.005. ns representing no significant difference. ** representing a significant

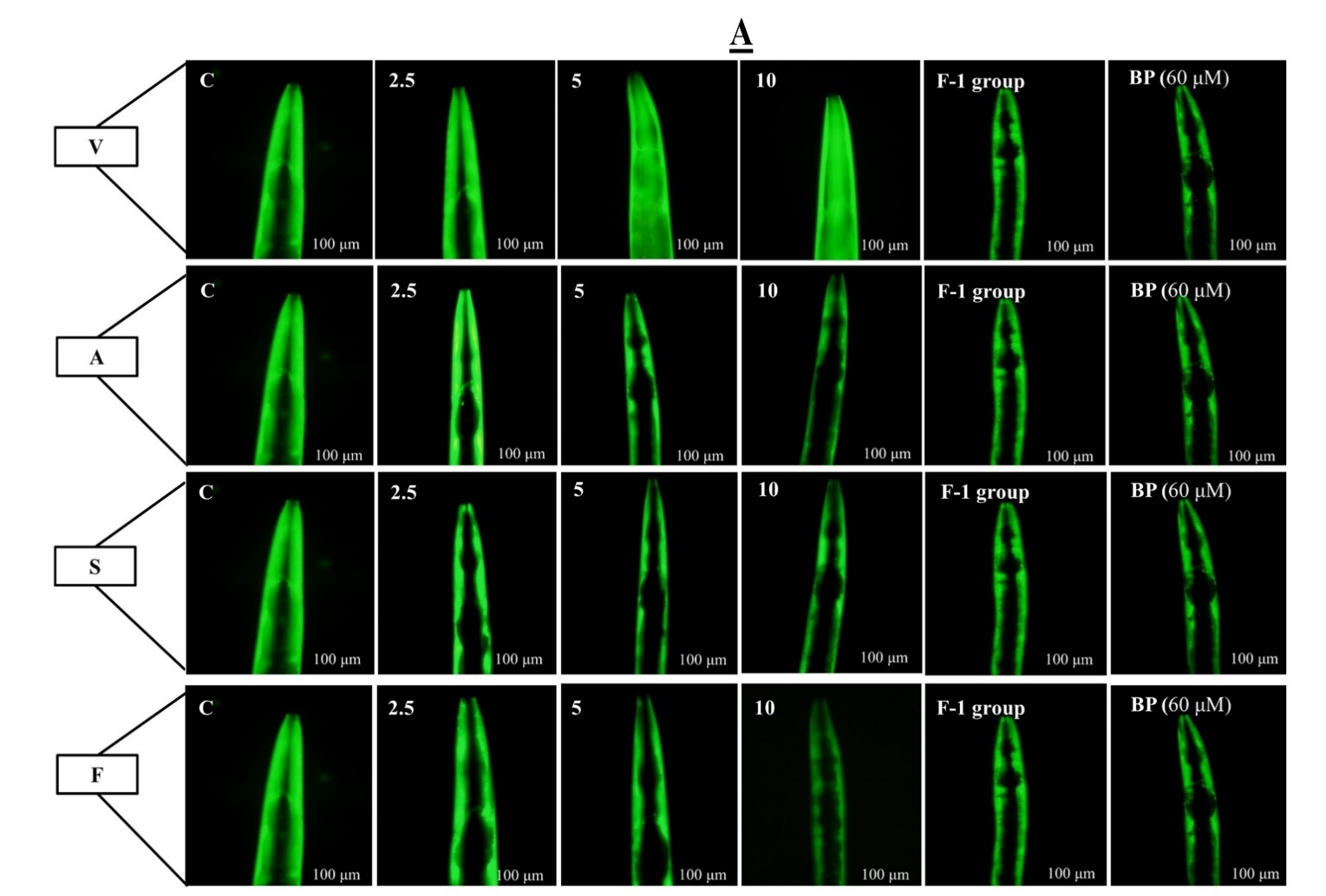

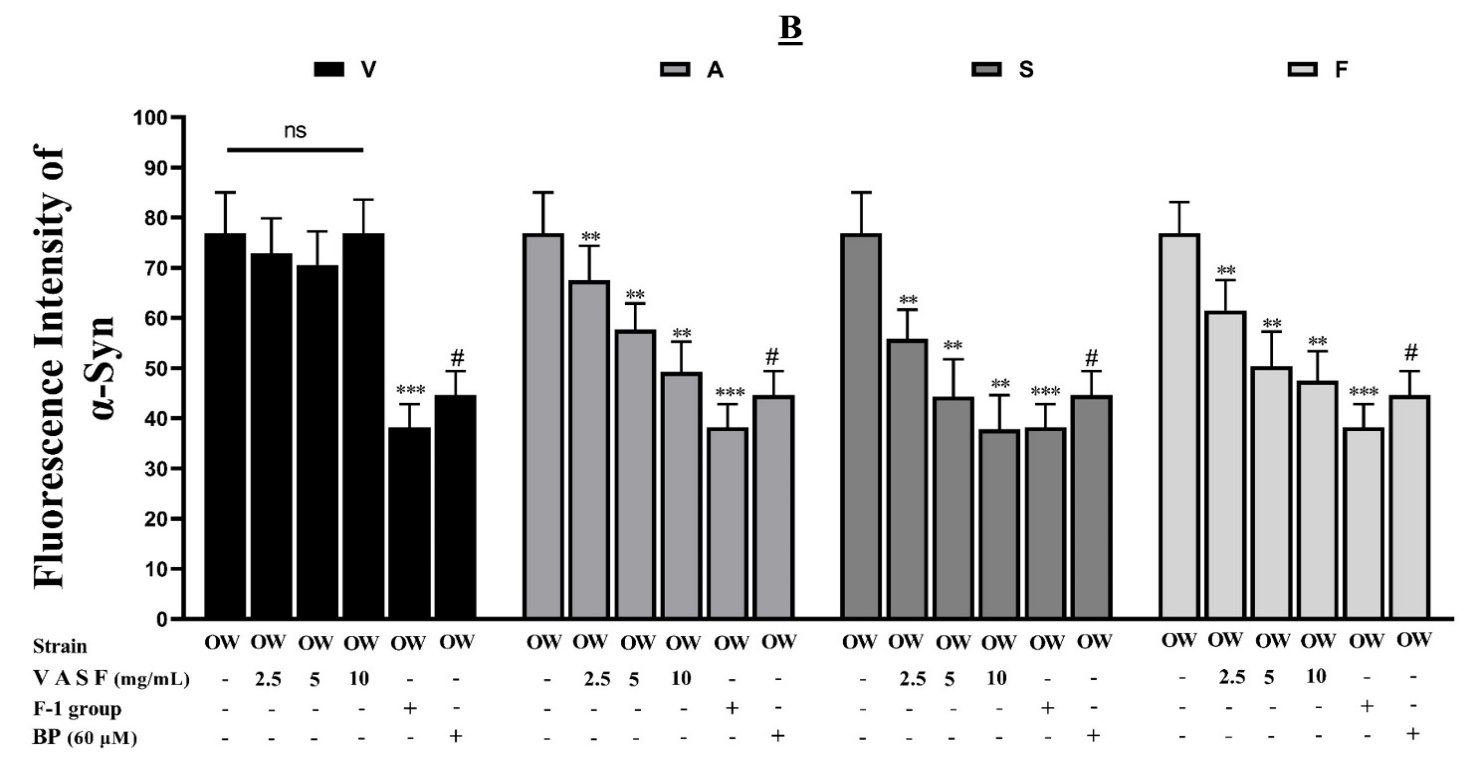
**(Supplementary figure 4A&4B).** Drug groups A, S, F, and F-1 on treatment reduced the α-Syn accumulated toxicity in transgenic worms OW13. **(A)** YFP fluorescence expressions in the muscles of worms OW13. The scale bar was 100 μm. **(B)** Bar Graph representing the quantitative data of the fluorescence intensities in OW13. ImageJ software was used to enumerate the fluorescence intensities of images. Data were computed by mean±SD (n = 40). ns depicting there is no significant difference between drug V dilutions on treatment. Similarly, ** shows a significant difference between the drugs A S F-treated and untreated groups (*p*≤0.005), and *** indicated a significant difference of *p≤*0.005 between the F-1 drug mixture and the untreated groups. While (#) indicates a significant difference of *p*≤0.005 between the positive control group BP (60 μM) with the untreated (control) groups on treatment.

of α-Syn toxicity (Bodhicharla et al., 2013). Initially, we treated the worms with VASF individually to analyze the effects of drugs via unc-54p: YFP expressions. The values were shown as the average intensity per body area. We found that A, S, F at 2.5 mg/mL, 5 mg/mL, 10 mg/mL in OW13 worms showed significant effects in reducing the α-Syn accumulated toxicity compared to untreated groups (Supplementary figure 4A). Graphical results in (Supplementary figure 4B) represent the fluorescent intensity calculated in OW13 worms treated with VASF or without VASF treatment at 2.5 mg/mL, 5 mg/mL, and 10 mg/mL. The fluorescent intensity of YFP expression in OW13, representing α-Syn protein expressions, significantly decreased by about 26% (A) *p*≤0.0005, 41 % (S) *p*≤0.005, and 34% (F) *p≤*0.005) compared to untreated groups. An S F drug group significantly reduced the α-Syn protein toxicity dose-dependently to drug group V (Supplementary figure 4B).

Next, we treated OW13 worms with the F-1 drug group. F-1 group assessment showed better results in decreasing the aggregative toxicity of α-Syn up to 40.01% *p*≤0.0005 than single drug-treated groups. Hence we can say that the drugs VASF in the decoction (F-1) supported and enhanced their protective actions against α-Syn accumulated toxicity much more effectively than the single drug group (Supplementary figure 4A-4B). BP (60 µM)) is used here as a positive control.

### 7 V, A, S, F and F-1 recovered Lipid Deposition in OW13 *C. elegans*

α-Syn is associated with fatty acid modifications and lipid content, which initiates vesicle formation through a specific mechanism (Lv et al., 2019). In OW13, the lipid contents were significantly reduced due to α-Syn expressions, disturbing its ubiquitin-like lipid compositions. Moreover, the toxic nature of accumulated α-Syn tends toward the rise of lipid peroxidation. It generates ROS, another leading cause of PD. We treated the OW13 with VASF at 2.5 mg/mL, 5 mg/mL, and 10 mg/mL and then used Nile red staining method to detect the lipid deposits. We
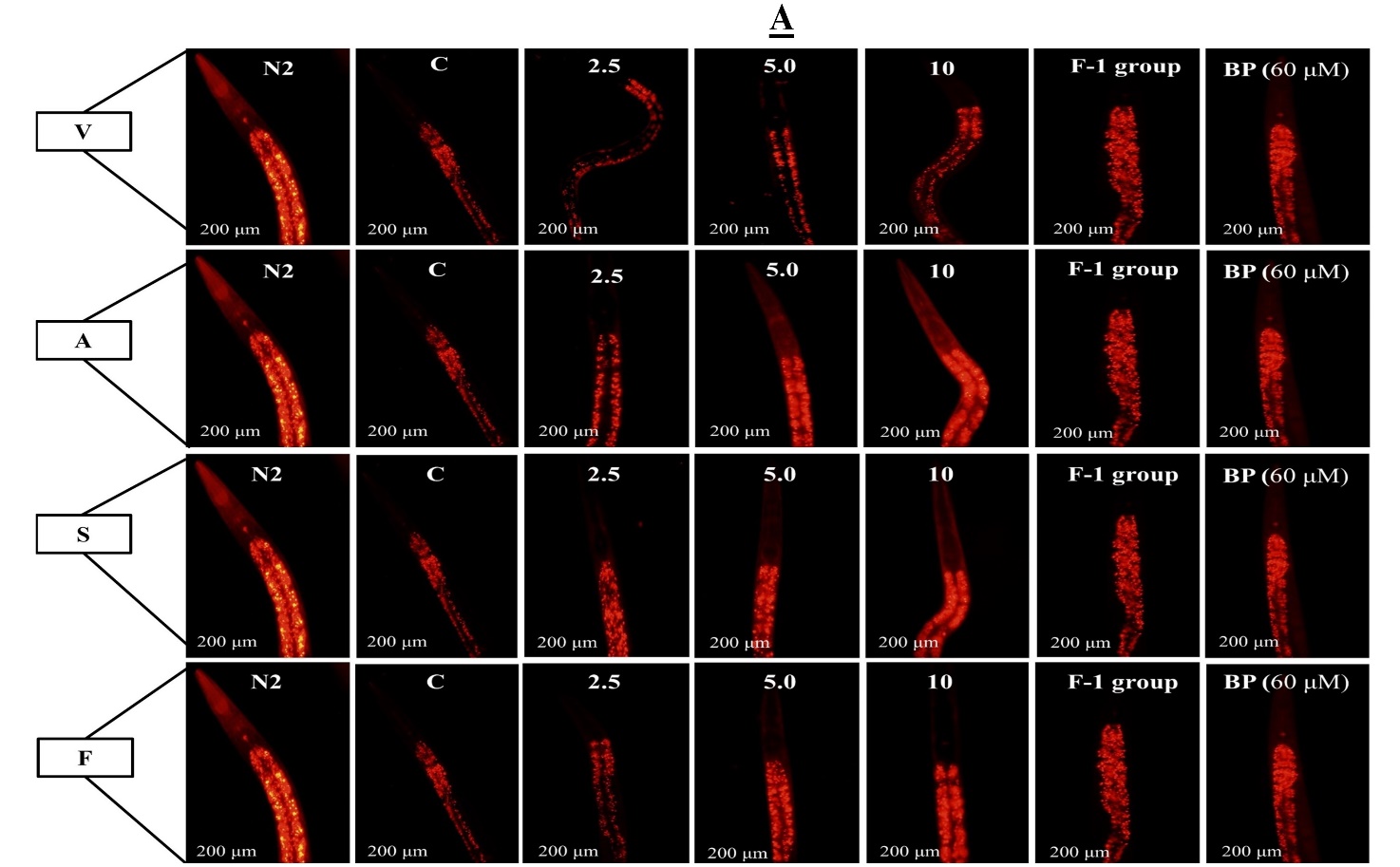

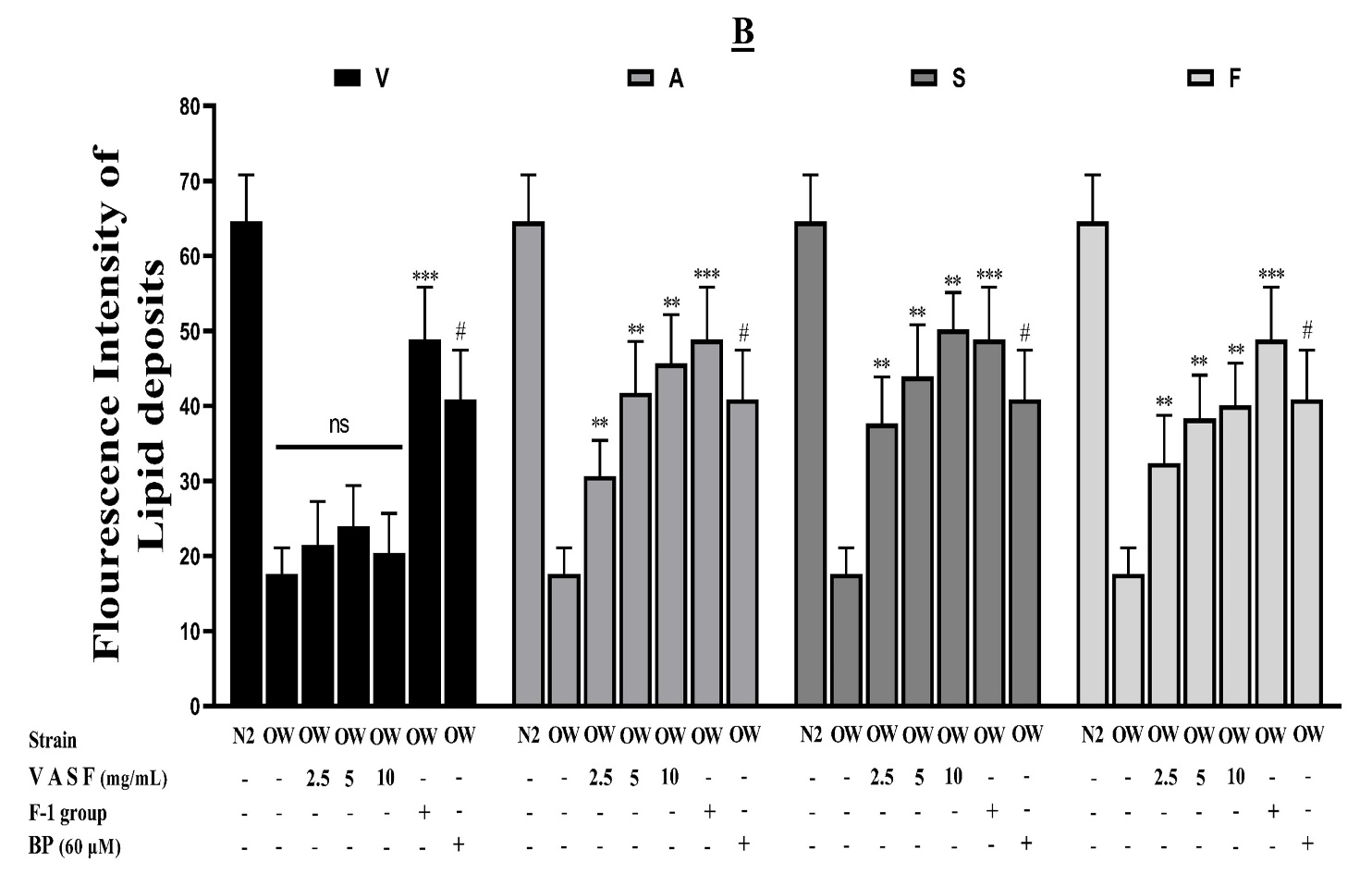

**(Supplementary figure 5A&5B).** Drug groups A S F and F-1 formula drug increased the lipid deposits in the transgenic worms OW13. **(A)** Representative images are the Nile red staining of OW13 worms after VASF treatment for 72 hrs. The scale bar was 200 μm. **(B)** A graphical picture of fluorescence intensities of strain OW13 was stained with the Nile red and quantified by software ImageJ. The data was calculated by mean ± SD (n = 30). ns depicting no significant difference compared with control between V drug dilutions. Similarly, drug group A S F showed a significant difference of *p*≤0.005 represented by ** between A S F treated and control groups. At the same time, *** describes the significant difference of *p*≤0.005 between the F-1 formula drugs group and the untreated groups. Here # represents the significant difference of *p*≤0.005 with the positive control **BP** (60 μM) and the untreated groups.

have conducted experiments to observe the lipid level in OW13 worms under both VASF treatment and non-treatment. Drug groups A S F significantly enhanced the lipid level dose-dependently on treatment except for the drug groups V **(A&B)**. Next, we treated the transgenic *C. elegans* with F-1 decoctions. Our results showed that F-1 group decoctions (formula) improved their anti-α-Syn actions and increased the lipid content in OW13 worms by about 24.331% more than the average of individual drugs on treatment. BO (60 µM) was used here as a positive control. *p*<0.005 between drug-treated and control groups.

**Table 1:** **Traditional Chinese Medicine with neuroprotective properties**

| Botanical  names | Abbr  (as) | Chinese names | Accepted Scientific  Names with History | Components | Traditional Uses | Ref |
| --- | --- | --- | --- | --- | --- | --- |
| Valeriana jatamansi | (V) | Zhi Zhu Xiang | *(*[nardostachyos radix et rhizoma](https://mpns.science.kew.org/mpns-portal/drugDetail?drugName=nardostachyos+radix+et+rhizoma&query=Valeriana+jatamansi+&filter=&fuzzy=false&nameType=all))  [Pharmacopoeia of China (2010)](https://mpns.science.kew.org/mpns-portal/reference?reference=100001&query=Valeriana+jatamansi+&filter=&fuzzy=false&nameType=all) [Pharmacopoeia of China (2015)](https://mpns.science.kew.org/mpns-portal/reference?reference=100050&query=Valeriana+jatamansi+&filter=&fuzzy=false&nameType=all) | Patchoul (24.3%)  α-bulnesene (13.8%)  Isovaleric acid (12.9%) | 1-Sedative  2-Anti-epilepsy  3-Nervous unrest | 14 |
| Acori Tatarinowii | (A) | Shichangpu | ([acori tatarinowii rhizoma](https://mpns.science.kew.org/mpns-portal/drugDetail?drugName=acori+tatarinowii+rhizoma&query=Rhizoma+Acori+Tatarinowii&filter=&fuzzy=false&nameType=all))  [Pharmacopoeia of China (2015)](https://mpns.science.kew.org/mpns-portal/reference?reference=100050&query=Rhizoma+Acori+Tatarinowii&filter=&fuzzy=false&nameType=all) | Α and β-asarone (11.3%, 9.41%)  (flavonoids) | 1-resuscitate  2-calm the mind,  3-resolve dampness | 65 |
| Fructus Schisandrae Chinensis | (F) | Wu Wei Zi | ([schisandrae chinensis fructus](https://mpns.science.kew.org/mpns-portal/drugDetail?drugName=schisandrae+chinensis+fructus&query=Fructus+Schisandrae&filter=&fuzzy=false&nameType=all)*)*  [Hong Kong Chinese Materia Med. Standards (2014)](https://mpns.science.kew.org/mpns-portal/reference?reference=8885&query=Fructus+Schisandrae&filter=&fuzzy=false&nameType=all) [Pharmacopoeia of China (2010)](https://mpns.science.kew.org/mpns-portal/reference?reference=100001&query=Fructus+Schisandrae&filter=&fuzzy=false&nameType=all) [Pharmacopoeia of China (2015)](https://mpns.science.kew.org/mpns-portal/reference?reference=100050&query=Fructus+Schisandrae&filter=&fuzzy=false&nameType=all) | Malic (27.98 mg/g) Citric (107.08 mg/g) Protocatechuic acids (0.42 mg/g) | 1-Liver disease  2-Relieve cough  3-Stomach disorder | 66 |
| Scutellaria baicalensis | (S) | HuangQin | ([scutellariae radix](https://mpns.science.kew.org/mpns-portal/drugDetail?drugName=scutellariae+radix&query=Scutellaria+baicalensis&filter=&fuzzy=false&nameType=all))  [Japanese Pharmacopoeia, 16th edn. (2012)](https://mpns.science.kew.org/mpns-portal/reference?reference=8870&query=Scutellaria+baicalensis&filter=&fuzzy=false&nameType=all) [Pharmacopoeia of China (2010)](https://mpns.science.kew.org/mpns-portal/reference?reference=100001&query=Scutellaria+baicalensis&filter=&fuzzy=false&nameType=all) [Pharmacopoeia of China (2015)](https://mpns.science.kew.org/mpns-portal/reference?reference=100050&query=Scutellaria+baicalensis&filter=&fuzzy=false&nameType=all) | Baicalein  (29.1%) | 1-Diarrhoea  2-Hypertension  3-Inflammation | 67 |

**Table 2: *C elegans* used in this study**

| **No** | **Strains** | **Transgene’s** | **Temp in ⁰C** | **Phenotypes** |
| --- | --- | --- | --- | --- |
| **1** | **N2** | Wild-type C. elegans | 20 | Wild type movement |
| **2** | **CL4176** | dvIs27 [myo-3p::*Aβ*_(1-42)_::let-851 3'UTR) + rol-6(su1006)] | 16 | Rapid paralysis |
| **3** | **CL2179** | myo-3/GFP, GFP control for CL4176 | 16 | Wt movement |
| **4** | **CL2355** | [snb1/*Aβ*_1-42_/long 3′-UTR + mtl-2::GFP]; | 16 | Hypersensitive to 5-HT |
| **5** | **CL2122** | dvIs15[(pPD30.38)unc-54(vector)+(pCL26)mtl 2::GFP];  control for CL2355 | 16 | Wt movement |
| **6** | **CL2006** | dvIs2 [pCL12 (unc-54/human *Aβ_1-42_* peptide minigene) + pRF4]; | 20 | Progressive paralysis |
| **7** | **TJ375** | [gpIs1(*hsp*-16.2::GFP)]; | 20 | Stress response analysis |
| **8** | **TJ356** | [zIs356 (Pdaf16::*daf*-16a/b::GFP+rol-6)]; | 20 | Reporter gene analysis |
| **9** | **CF1553** | [muIs84 ((pAD76) *sod*-3p::GFP +rol-6)]; | 20 | Antioxidative enzymes expression |

**Table 3: Defining the V, A, F, S drug Orthogonal Experiments for Evaluating the F-1 formula Percentage**

| **Groups and Concentrations of V, A, F, S TCM for orthogonal experiments** | | | | |
| --- | --- | --- | --- | --- |
| **Groups/Cons** | **S (mg/mL)** | **F (mg/mL)** | **A (mg/mL)** | **V (mg/mL)** |
| Group 1 | 0 | 0 | 0 | 0 |
| Group 2 | 0 | 2.5 | 10 | 10 |
| Group 3 | 0 | 5.0 | 20 | 20 |
| Group 4 | 2.5 | 0 | 10 | 20 |
| Group 5 | 2.5 | 2.5 | 20 | 0 |
| Group 6 | 2.5 | 5.0 | 0 | 10 |
| Group 7 | 5.0 | 0 | 20 | 10 |
| Group 8 | 5.0 | 2.5 | 0 | 20 |
| Group 9 | 5.0 | 5.0 | 10 | 0 |
